# Common rules underlying optogenetic and behavioral modulation of responses in multi-cell-type V1 circuits

**DOI:** 10.1101/2020.11.11.378729

**Authors:** Agostina Palmigiano, Francesco Fumarola, Daniel P. Mossing, Nataliya Kraynyukova, Hillel Adesnik, Kenneth D. Miller

## Abstract

The visual cortex receives non-sensory inputs containing behavioral and brain state information. Here we propose a parallel between optogenetic and behavioral modulations of activity and characterize their impact on cell-type-specific V1 processing under a common theoretical framework. We infer cell-type-specific circuitry from large-scale V1 recordings and demonstrate that, given strong recurrent excitation, the cell-type-specific responses imply key aspects of the known connectivity. In the inferred models, parvalbumin-expressing (PV), but not other, interneurons have responses to perturbations that we show theoretically imply that their activity stabilizes the circuit. We infer inputs that explain locomotion-induced changes in firing rates and find that, contrary to hypotheses of simple disinhibition, locomotory drive to VIP cells and to SOM cells largely cancel, with enhancement of excitatory-cell visual responses likely due to direct locomotory drive to them. We show that this SOM/VIP cancellation is a property emerging from V1 connectivity structure.

## Introduction

The visual cortex of the mouse represents a variety of different signals beyond visual information. Eye position (Meyer et al., 2018; Parker et al., 2022), face (Stringer et al., 2019) and head (Bouvier et al., 2020; Vélez-Fort et al., 2018) movements, and locomotion (Ayaz et al., 2013; Dipoppa et al., 2018; Fu et al., 2014; Keller et al., 2012; Niell and Stryker, 2010) are represented together with visual information in the cortical activity of mouse V1. The integration of locomotory and visual signals is at least partially mediated by the actions of specific interneuronal cell types (Dipoppa et al., 2018; Fu et al., 2014). Three types of GABA-ergic cells – parvalbumin-(PV), somatostatin-(SOM), and vasoactive-intestinal-peptide-(VIP) expressing cells – form a microcircuit characterized by highly specific connectivity (Campagnola et al., 2022; Karnani et al., 2016; Pfeffer et al., 2013), that can simultaneously stabilize and loosely balance strong recurrent excitation (Adesnik, 2017; Ahmadian and Miller, 2021; Ozeki et al., 2009), gate top-down signals (Zhang et al., 2014, 2016), and control the gain of the pyramidal cell population (Pi et al., 2013). In particular, two of these types, SOM and VIP, engage in competitive dynamics whose outcome directly regulates pyramidal cell activity and underlies modulation by contextual (Adesnik et al., 2012; Keller et al., 2020a,b; Mossing et al., 2021; Schnabel et al., 2018; Veit et al., 2017) and behavioral (Ayaz et al., 2013; Dipoppa et al., 2018; Fu et al., 2014; Niell and Stryker, 2010) state. Nevertheless, the mechanisms used by this circuit to simultaneously implement the multiple computations it performs, the dynamical signatures of those computations, and their linkage to the circuit’s connectivity are incompletely understood.

Here we propose that optogenetic and behavioral modulations of activity may share common features, and describe them by a common perturbative framework. First, we develop a program for inferring from data recurrent circuit models that capture the full distribution across neurons of each cell type’s responses. We apply this framework to calcium recordings of responses to visual stimuli of pyramidal (E) cells and the above three inhibitory cell types in mouse V1 layers 2/3, in stationary (non-locomoting) mice, and infer statistical descriptions of the recurrent circuitry that capture the distributions of responses of these cell types. The models we infer from data exhibit key aspects of the structure of the connectivity found in the mouse visual system – namely weak or no connections from VIP to any cell type except SOM or from SOM to itself (Karnani et al., 2016; Pfeffer et al., 2013) – provided that the recurrent excitation is strong, suggesting that this structure is needed to generate the observed responses given strong recurrent excitation. Second, we develop a theoretical framework for quantitatively predicting the distributions of responses of each cell type in model circuits to patterned, full or partial optogenetic perturbations. We show that if the mean response of a given cell-type population to a perturbation of that cell type is “paradoxical” – a negative response to positive stimulation – then that population is stabilizing the circuit. We predict that PV but not SOM interneurons show such “paradoxical” responses, and do so across stimulus conditions. This is consistent with convergent experimental (Veit et al., 2017) and theoretical (Bos et al., 2020) arguments that PV interneurons, but not SOM interneurons, play a major role in circuit stabilization. Furthermore, the fraction of PV cells that respond paradoxically is found to depend non-monotonically on the fraction of PV cells that are stimulated, with the turnaround point determined by the fraction of PV cells that are necessary to stabilize the circuit. This *fractional paradoxical effect* opens a new avenue for experimental inquiry.

Finally, under the hypothesis that locomotion acts by adding inputs of different strengths to the different cell types in our inferred circuits, we leverage our theoretical framework to infer the locomotion-induced inputs that can account for the cell-type specific distributions of response changes observed with locomotion. We find that these response changes require that locomotion induces strong inputs to both VIP and to SOM, whose effects on non-VIP cells largely cancel. This cancellation explains the lack of locomotory response of E cells in the absence of visual stimulation, despite the disinhibitory effect expected from the locomotory activation of VIP cells (Dipoppa et al., 2018; Pakan et al., 2016). We show that this cancellation of the effects of VIP and SOM stimulation is a general property of the inferred circuit structure, and is reliably seen at the level of single-cell responses as well as at the population level. Our results suggest that locomotion-induced increases in pyramidal cell activity in the presence of visual inputs is induced by a locomotion-induced increase in visual drive to E cells, consistent with the fact that acetylcholine release, which is induced by locomotion (Fu et al., 2014), increases the gain of thalamocortical synapses (Gil et al., 1997; Hsieh et al., 2000). This is a first example of a more general approach to inferring inputs underlying behavioral modulations of neural responses.

## Results

### Inference of large scale models from multi-cell-type data recover V1 connectivity structure

In order to investigate the mechanisms that underlie responses to behavioral and optogenetic perturbations in mouse V1, we build large-scale, cell-type-specific models by inferring the model parameters from data. We consider the response to visual stimuli of varying contrasts in neurons of layer 2/3 of primary visual cortex (V1) of awake, head-fixed mice. Specifically, we study the responses, as assayed by calcium imaging, of Pyramidal (E) cells and of Parvalbumin (PV), Somatostatin (SOM) and Vasoactive Intestinal Polypeptide (VIP) expressing interneurons while the animal is shown square patches of drifting grating stimuli of a small size (5 degrees) at varying contrasts.

Our large-scale model is a fully recurrent network composed of four populations representing the four cell types, with many units of each cell type (Fig. 1**a**). Each unit has a rectified power-law input-output function (Ahmadian et al., 2013) and receives a baseline input to account for the spontaneous activity, while feed-forward inputs carrying visual information only target E and PV cells. In order to find parameters for this model, we begin by first inferring the parameters of a simplified recurrent circuit model with only four units, one unit per population, each representing the mean activity of its cell type’s population in the case in which there is no heterogeneity among the cells within each population (see Fig. 1**b**, fitting pipeline). From the measured means and standard errors of the responses of each cell type to each contrast, we sample millions of sets of surrogate curves of mean response vs. contrast for the four cell types, and use non-negative least squares to fit a model to each surrogate set (following Renner et al. (2020), see Fig. S1 and Methods section 3.2). We then select, from these millions of possible models, some hundreds of data-compatible 4-unit models for which the network steady-states provide the best fit.

**Figure 1.**
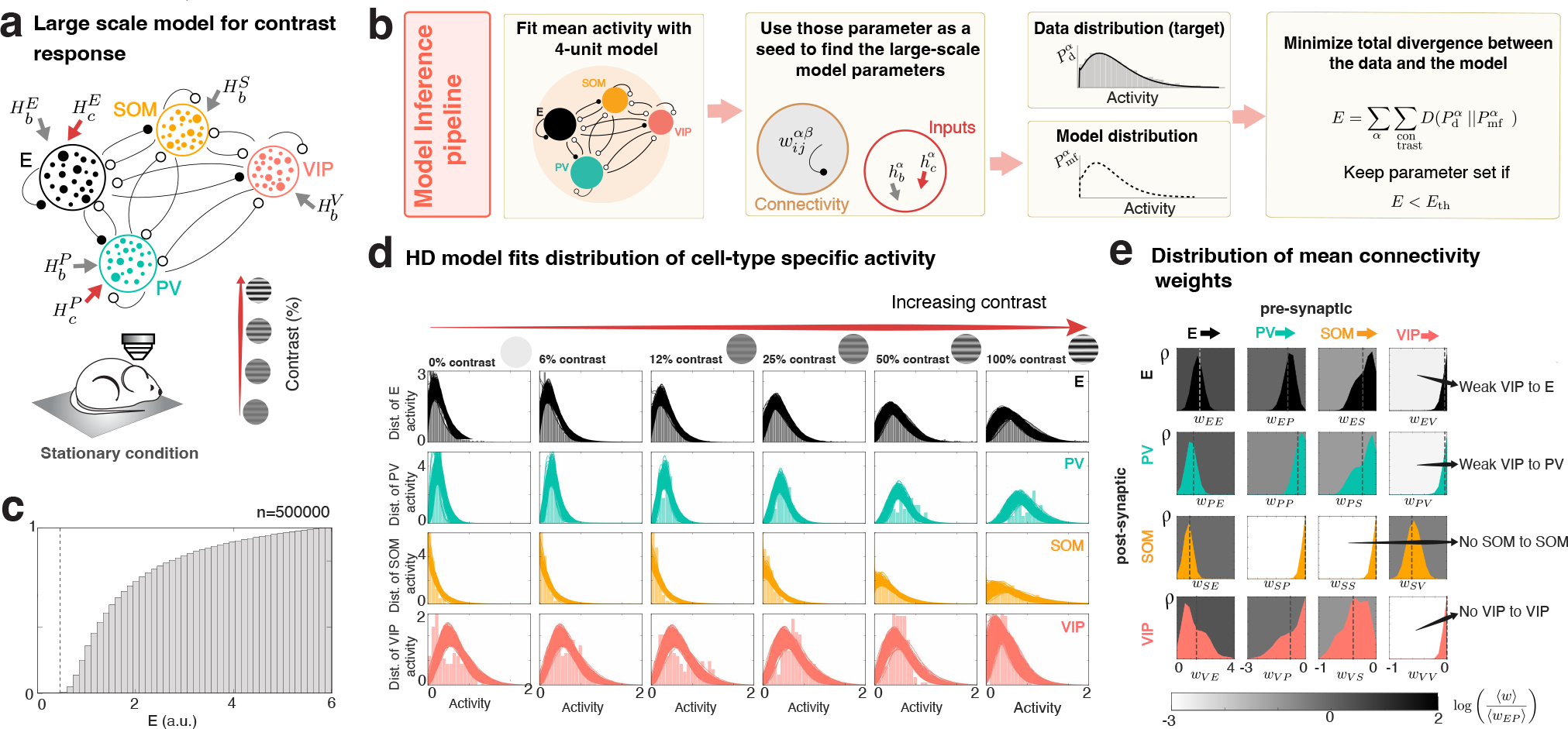
Large scale model of mouse V1. **a)** Large scale model composed of pyramidal (E, black), PV (turquoise), SOM (orange) and VIP (pink) populations. Inputs to each cell type are composed of a spontaneous activity baseline *H_b_* and a stimulus related current, *H_c_* modeling the feed-forward inputs from layer 4 targeting only E and PV (see also Fig. S1). **b)** Inference pipeline. Left: Model of four populations (see also (Renner et al., 2020)) describing only the mean activity. This model is fitted to the mean activity of the data (see Fig. S1 for details). Middle and middle right: we generate large-scale models by sampling from prior distributions of parameters given by the 4 unit model fits, computing the distributions of responses produced by the large-scale model with those parameters and finally comparing the model distributions we obtain, to the data distributions using an error function. Right: The error is given by the sum of the Kullback-Leibler divergences of the distributions given by the model (*P*_mf_) and the data (*P*_d_) for all cell types and contrast values, which can be found explicitly. We build a suitable family of models by only accepting errors smaller than a threshold of 0.5 (top 0.005%) (in other words we perform approximate Bayesian computation (Sisson et al., 2018)). **c)** Distribution of KL divergences, indicating the 0.5 threshold. We used models below this threshold for the analysis in the remaining text (see Methods section 4.4 for details). **d)** Data (histogram, colored bars) and data fits (solid colored line) are in good agreement. **e)** Distribution of mean connectivity weights over all possible models is shown. The gray-scale background of each panel is the logarithm of the absolute value of the mean of each distribution. As in experiments (see Fig.S1), the models are found to have, at most, very weak recurrent SOM and VIP connections and connections from VIP to E and PV.

We then fit the large-scale model. Its parameters are the means and variances of the baseline inputs to all cell types, of the contrast-driven inputs to E and PV, and of the connectivities from each cell type to itself and to each of the other cell types. To fit this model, we use the parameters of the hundreds of fitted 4-unit models as initial, or *seed* parameters for the means, and search over priors to find the variances (see Methods section 4.4). To fit this model, we leverage two facts. First, in large-scale circuit models in which neurons have a power-law input-output function, there is an explicit expression describing the distributions of activity that the population of each cell type produces (Roxin et al., 2011) given the distribution of its total inputs (recurrent and external). Second, for a fixed set of parameters, the distributions of activities the model will produce can be computed self-consistently through mean-field theory (Cabana and Touboul, 2013; Kadmon and Sompolinsky, 2015). From these two sets of analytic expressions, we create an explicit error function that quantifies how different the measured distributions of activity of all cell types at all contrasts are from those distributions produced by the large-scale model with a given set of parameters (see Methods sections 4 and 4.4 for a full description of the method). We construct a family of large scale models that match well the experimentally measured distributions of responses of all of the cell types (see below and Fig. S1) by choosing those that minimize that error function (Fig. 1**c-d**). This family of models recapitulates the dependence of the distribution of responses of all cell types on contrast, and captures both the spreading out of the distributions with increasing contrast and the tails of the distributions seen in calcium data. These results do not depend on the exact shape of the distribution of weights as long as these distributions are bounded. Remarkably, even models with log-normally distributed synaptic weights reproduce the data similarly well when using the exact same statistics as the models shown here (Fig. S1).

We required the model to have strong recurrent excitation (strong enough to be an inhibition-stabilized network, or ISN, described further below, see Fig. S1**c-d**). Beyond that, this optimization takes as sole input the response data and uses no prior information on the synaptic structure. It is not obvious that any meaningful synaptic structure should be recoverable from such a procedure. Yet, across models that fit the response distributions, the structure of the inferred connectivity matrices has a striking resemblance to that reported experimentally (Fig. 1**e**, see also S1**b**). In particular, recurrent connections within the SOM population and the VIP population were weak or absent, and VIP interneurons had weak or absent connections to all other cell types except SOM interneurons; these are both robustly reported features of the connectivity of mouse V1 L2/3 (Campagnola et al., 2022; Karnani et al., 2016; Pfeffer et al., 2013).

### PV stabilizes the cortical circuit and is the only cell-type with paradoxical response

Cortical circuits have strong recurrent excitation that is stabilized and loosely balanced by inhibition (Adesnik, 2017; Ahmadian and Miller, 2021; Kato et al., 2017; Moore et al., 2018; Ozeki et al., 2009; Sanzeni et al., 2020). In models of one excitatory and only one inhibitory population, inhibitory stabilization implies that an increase in the input drive to the inhibitory population will elicit a “paradoxical” *decrease* of the inhibitory steady-state activity, along with a decrease of the excitatory activity (Ozeki et al., 2009; Tsodyks et al., 1997). In models with multiple cell types, if a perturbation only to GABAergic cells elicits a decrease in excitatory-cell activity along with a paradoxical decrease in the inhibition received by E cells – the latter would necessarily be true if there was a decrease in the activity of all GABAergic cells that project to excitatory cells – then the circuit is an ISN (Litwin-Kumar et al., 2016; Rubin et al., 2015). The converse, that the ISN condition implies a paradoxical response of the inhibitory activity, is only true in an E/I circuit: in the multi-cell-type case, there are multiple ways in which the total inhibitory input current to the E population can decrease, so no specific cell type needs to decrease its activity.

What then are the conditions that need to be satisfied for a set of cells to respond paradoxically to stimulation? Answering this question will lead us to an understanding of how stabilization of strong recurrent excitation is distributed across the different inhibitory cell types. We show theoretically, that paradoxical response of a given cell-type population implies that the circuit without that population is unstable (see Fig. 2). In other words, if a cell-type responds paradoxically, then if we would clamp the activity of that cell-type, not allowing it to change to track the activity of the rest of the circuit, a small perturbation to the circuit would make the activity grow or collapse unchecked. Furthermore, we find that because of the structure of the connectivity, a paradoxical response of SOM directly implies that the E-PV sub-circuit is unstable. A summary of the results derived in this section are: we find that i) the mean response of PV to its own stimulation is paradoxical in almost all the models that fit the data across stimulus conditions, and that ii) almost all models without PV are unstable, while iii) SOM and VIP don’t have a mean paradoxical response, and iv) sub-circuits without SOM and/or VIP are always stable. These results are consistent with experimental work showing that strong perturbations to PV destabilize the dynamics in V1 while strong perturbations to SOM do not (Veit et al., 2017).

**Figure 2.**
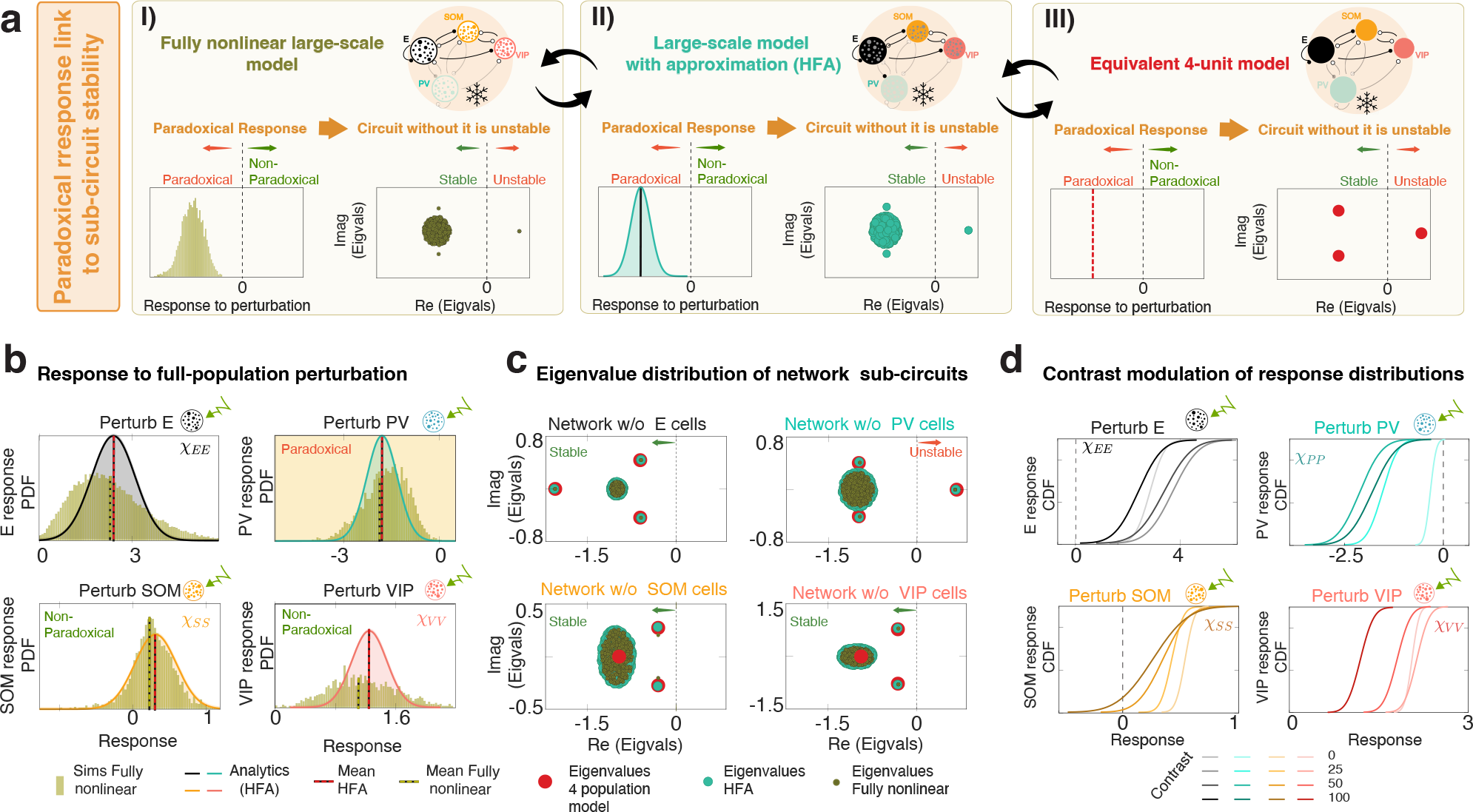
Paradoxical response and circuit stabilization of inferred models. **a)** Graphic summary of the relation between stability and paradoxical responses. (I): We observe a paradoxical PV response in the fully nonlinear system seems to imply instability of the network without that cell type (one eigenvalue has positive real part). (II) We find that under the HFA (see text), the distribution of responses has mean (red line) given by the response of the 4-unit model, and a variance that can be computed exactly. Furthermore, the outlier eigenvalues in this case are given by the 4-unit model. (III) In the 4-unit model, a negative (paradoxical) response of a cell type to a positive perturbation (red dashed line, left) provably implies that the circuit without that cell type is unstable (see Eq. 1), which means that at least one eigenvalue of the Jacobian of the circuit without that cell type has a positive real part (right). **b)** Distribution of responses of each cell type to its own full-population perturbations for a stimulus of 100% contrast, for the fully nonlinear large-scale system (green, dashed green line is its mean), and the analytical distributions obtained under the HFA (colored histograms, red dashed line is its mean). The responses of E, VIP, and most SOM cells are not paradoxical, while all cells in the PV population respond paradoxically to PV stimulation. When VIP projects only to SOM, which is true in most our models, the lack of a paradoxical response in SOM indicates that the E-PV circuit is stable. **c)** Eigenvalue distribution of the Jacobian of the sub-system without the E (top left), without the PV (top right), without the SOM (bottom left) and without the VIP (bottom right) populations. Only the sub-circuit without the PV population is unstable (has a positive eigenvalue). The eigenvalues of the fully nonlinear system overlap with those of the system under the HFA, which coincide with those of the 4-unit model. **d)** Cumulative distribution of responses (HFA) to full-population perturbation in the presence of a visual stimulus for varying stimulus contrast. These results hold in the family of models that fit the data, see Fig. S2.

In order to demonstrate these results theoretically, we establish a formalism that allows us to study response to (small) perturbations of arbitrary fractions of cells of each cell type, and how the response is related to the stability of the network without that cell type, in large-scale models (Fig. 2**a**(I)). The theory relies on a simplifying approximation, which is that the gain (the slope of the input/output function) of each cell is the same for all cells in a given cell-type population. Under this approximation, which we refer to as the *homogeneous fixed point approximation* (hereinafter HFA, see Methods section 5.2), and by building on recent random matrix theory results (Ahmadian et al., 2015) (see Methods section 5), we are able to obtain explicit expressions for the behavior of the mean (See. Eq. S84 and section 5.5 of the Methods) and the variance (Eq.S108 in section 5.6 of the Methods) of the distributions of responses in each population to either full or partial, and either homogeneous or heterogeneous, perturbations. Importantly, we find that under the HFA: i) the mean responses to a perturbation of all neurons of a given cell type are equal to the responses in a corresponding 4-unit model to a perturbation of that cell type, and ii) when that mean response is paradoxical, the sub-circuit without that cell type is unstable (Fig. 2a(III); see Methods sections 5.11, 6). Thus, we can use the HFA as an analytic bridge between the simple 4-unit model and the fully nonlinear large-scale model.

To state these results mathematically, we first consider the steady-state responses to a small perturbation of a 4-unit circuit, initially at a fixed point (a steady-state response to a fixed input). The responses are described by the linear response matrix *χ*. The response of cell type *α* to a perturbative input to itself is given by the diagonal entry *χ_αα_* multiplied by the input. When *χ_αα_* is negative, this response is paradoxical – a negative response to a positive input. *χ_αα_* in turn can be written as a function of the Jacobian, *J_α_*, of the sub-circuit without the cell-type *α*, at the fixed point (see also Methods sections 3.3 and 3.3.2):

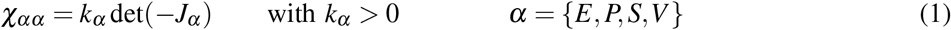

The Jacobian determines the stability of the sub-circuit. For a fixed point of a system to be stable, all eigenvalues of the negative Jacobian have to have positive real part (corresponding to all eigenvalues of the Jacobian having negative real part), and thus its determinant – the product of its eigenvalues – must be positive. Thus, if the response of cell type *α* at a given fixed point is paradoxical (*χ_αα_* < 0), then, at that fixed point, the sub-circuit without that cell type is unstable (det(*J_α_*) < 0), *i.e*. at least one eigenvalue of the Jacobian of the sub-circuit has positive real part (Fig. 2a). This insight is a simple generalization of the two-population ISN network – there, when the sub-circuit without I cells was unstable, then I cells responded paradoxically – that links cell-type specific paradoxical response to sub-circuit stability more generally (Fig. 2**a**, left). We furthermore find that when VIP projects only to SOM (as is at least approximately true in mouse V1), then a paradoxical response in the SOM population indicates not only that the E-PV-VIP sub-circuit is unstable, but that the E-PV sub-circuit by itself is unstable (see Eq. S14). These simple findings in the model with 4 units apply more generally: we find that in the large scale model, under the HFA, whenever the mean response of a cell-type to a perturbation of the entire population is paradoxical, the system without that population will be unstable (Fig. 2**a**, middle, see theoretical argument for this in Methods section 6), and we have found this to hold in numerical simulations of the fully nonlinear system (Fig. 2**a**, right).

We find (as illustrated for an example model at 100% contrast, Figs. 2b-c) that in all three models (the 4-unit model, the large-scale model with the HFA, and the fully nonlinear large-scale model) the mean response is only paradoxical (negative) for PV cells, and that therefore the system without PV is unstable (it has one positive eigenvalue, top-right panel). The PV responses remain paradoxical, and that of the other cell types non-paradoxical, at all stimulus contrasts (Figure 2**d**). These results hold in almost all the models that fit the data (see S2**a-b**), suggesting a fundamental role of PV in circuit stabilization in the family of models we derived from the data. Furthermore, we find that, in the great majority of solutions, no other interneuron type has a mean paradoxical response, and that sub-circuits without them (*i.e*., without SOM and/or VIP) are always stable (Fig. S2**a-d**).

In summary, we find that in most models that fit the data i) SOM does not respond paradoxically, consistent with the E-PV circuit being stable, and ii) PV responds paradoxically, meaning that the circuit without it is unstable (Fig. 2**b,c**). This is consistent with experimental work showing that suppression of PV activity destabilizes V1 while suppression of SOM activity does not (Veit et al., 2017)

### Fractional paradoxical effect

Motivated by the fact that in optogenetic experiments with viral injection only a fraction of cells is perturbed, we next investigated how many PV cells need to be stimulated to observe the paradoxical effect in the population of PV cells (Fig. 3). We find, that, when targeting only a fraction of the cells in the circuit, there is a range in which the more strongly we probe the system to find the paradoxical effect, the less likely we are to find it. Specifically, we find that there is a range in which the more PV cells we perturb (which experimentally can be done in a controlled manner via holographic optogenetics, Adesnik and Abdeladim, 2021) the less likely we are to observe a paradoxical effect by measuring the activity of the population. We name this *the fractional paradoxical effect*. Furthermore we find, that under the HFA, the perturbed cells only have a mean paradoxical response when the sub-circuit without the perturbed cells becomes unstable, generalizing the above findings to partial perturbations.

**Figure 3.**
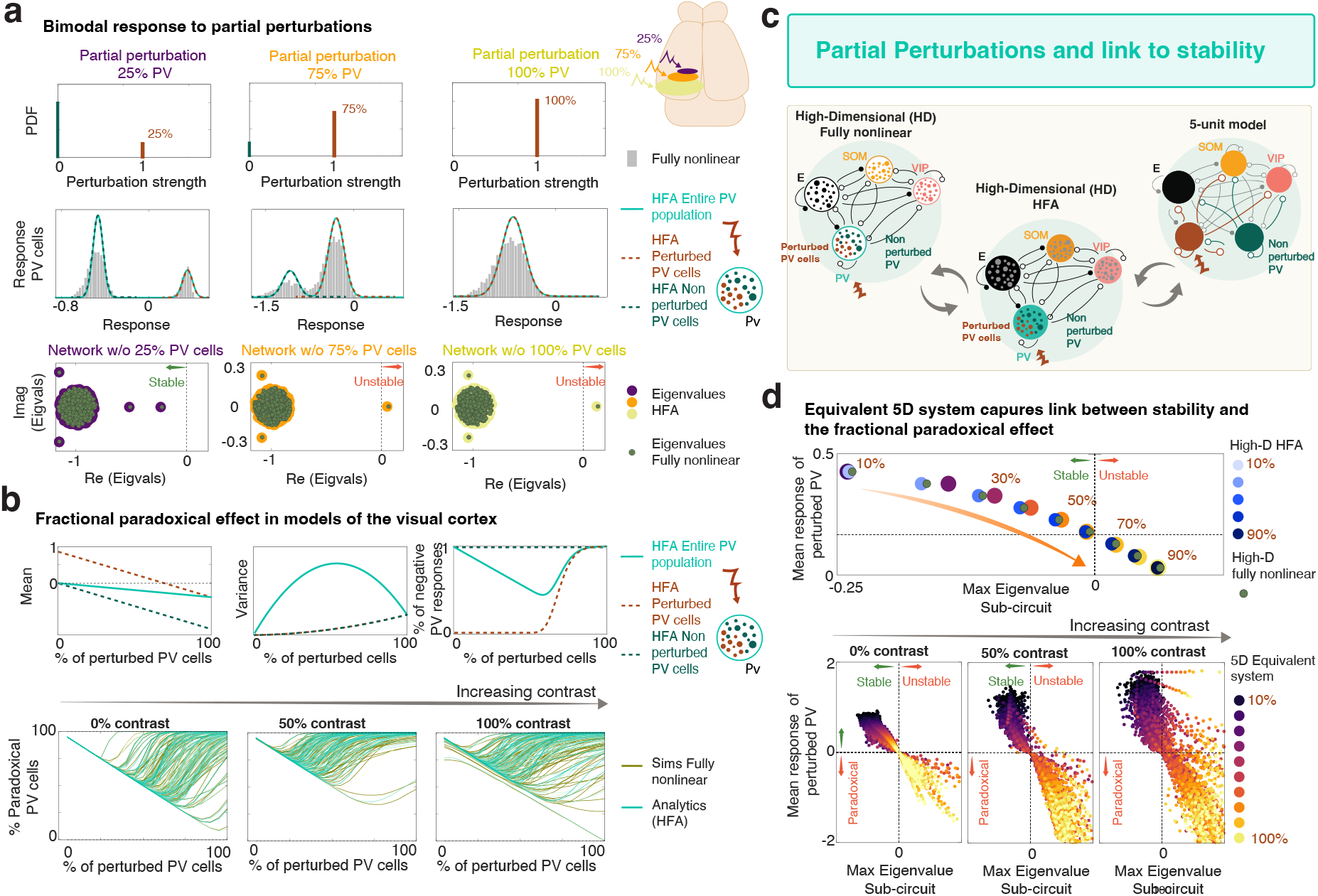
Fractional paradoxical effect and link to sub-circuit stability: **a)** Top: distribution of perturbation strengths, when perturbing 25% (left), 75% (middle) and a 100% (right) of the PV population. Middle row: Partial perturbations result in a bimodal distribution of responses. Simulations (gray) are well described by the mixture of two Gaussians predicted by the HFA (turquoise, solid line). The rightmost peak (dashed red) corresponds to the response of the sub-population of stimulated PV cells, while the leftmost peak (dashed green) corresponds to the response of non-stimulated PV cells. Note that the mean response of the perturbed population changes sign with increasing number of perturbed PV cells. Bottom: Eigenvalue spectrum of the Jacobian of the non-perturbed sub-circuit for the HFA approximation (purple, orange and yellow) and the fully nonlinear system (green). The maximum eigenvalue of the sub-circuit changes sign, indicating that the non-perturbed sub-circuit has become unstable, with increasing number of perturbed PV cells. **b)** Top left: Mean of the entire (bimodal) distribution of PV cell responses (turquoise), the mean of the perturbed PV cell responses (dashed red) and the non-perturbed PV cell responses (dashed green) as a function of the fraction of PV cells perturbed. Top middle: While all three means monotonically decrease with the fraction of stimulated cells, the variance of both non-perturbed (dashed green) and perturbed (dashed red) monotonically increase, while the distance between the two peaks, and thus the variance associated with this distance, monotonically decreases (not shown), resulting in a non-monotonic change in variance of the full distribution (turquoise). Top right: Fraction of negative responses as a function of the fraction of stimulated cells shows a non-monotonic dependence (turquoise), which we name the *fractional paradoxical* effect. Bottom: The fractional paradoxical effect is a signature of models that fit the data, and occurs for all values of the contrast. Simulations of the fully nonlinear system for different data-fit models (green) are in good agreement with calculations from the HFA for the same models (turquoise). **c)** Linking response and stability across models. A fully nonlinear system can be linked to a large-scale system of lower complexity via the HFA. The mean response to a partial perturbation in the HFA can be mapped to the response to perturbation of one PV sub-population in a 5-unit system with two PV sub-populations, a perturbed one (red) and an unperturbed one (green) see, Eq. (S124). **d)** Top: Mean response of the perturbed PV population as a function of the outlier eigenvalue with largest real part of the Jacobian of the unperturbed subcircuit, for different fractions of perturbed PV cells (numbers in graph indicate this fraction). This is shown for the fully nonlinear system (green), HFA (blue colors) and the equivalent 5D system (purple/orange palette, legend in bottom panel). The mean responses become negative when the maximum eigenvalue real part crosses zero, indicating instability of the non-perturbed sub-circuit. Bottom: Mean response of the perturbed PV population, for varying fractions of perturbed cells (indicated by colors), as a function of the value of the outlier eigenvalue in the equivalent 5D model obtained from different models that fit the data, for three different values of the contrast (see also Methods section 5.9).

Mathematically, we establish these results as follows. Under the HFA, the distribution of responses of the entire population to partial perturbations is bimodal, given by a mixture of two Gaussians (Fig. 3**a**, turquoise). One Gaussian distribution corresponds to the perturbed cells (red dashed line), while the other corresponds to the unperturbed population (green dashed line, see Eq. S113).For the PV population, the unperturbed cells always show a negative mean response to the perturbation. When the number of perturbed PV cells is small, the mean of the Gaussian response distribution corresponding to the perturbed cells is positive (see also Fig. 3**b**) and the sub-circuit without those perturbed cells is stable (all the eigenvalues of the sub-circuit’s Jacobian have negative real part (Fig. 3**a**, bottom left; Fig. S3). As the fraction of perturbed PV cells increases, the mean response of the perturbed population moves towards negative values. However, since the fraction of perturbed PV cells is increasing, then while their response remains positive, the fraction of negatively responding PV cells is decreasing. With continued growth of the perturbed fraction, their response ultimately changes sign, as the sub-circuit without the perturbed cells loses stability (the maximum eigenvalue of the Jacobian of the sub-circuit without the perturbed cells becomes positive, Fig. 3**a**, bottom right). This gives rise to a curious phenomenon: with increasing fraction of PV cells perturbed, the fraction of PV cells responding negatively shows non-monotonic behavior (Fig. 3**b**, top right). Over some range, increasing the fraction of stimulated PV cells decreases the probability that we will measure a PV cell showing negative response. This *fractional paradoxical effect* extends the concept of critical fraction (the fraction of inhibitory neurons that must be perturbed in order to see a paradoxical effect, developed by Sadeh et al. (2017)), to the case in which the neurons have heterogeneous connectivity. The lower panels of Figure 3**b** show the dependence of the fraction of PV negative responses on the fraction of perturbed PV cells for different values of the stimulus contrast in the models obtained in Figure 1. Intriguingly, in the models that fit the data, PV has a fractional paradoxical response at all contrasts. Recent experiments (Sanzeni et al., 2020) have revealed that an optogenetic perturbation of PV interneurons with transgenic opsin expression (affecting essentially all PV cells) elicits a paradoxical effect in most cells, whereas if the expression is viral (and therefore affects only a fraction of PV cells), a much smaller portion (about 50%) of cells show negative responses. Our models are consistent with that observation, and predict that this property is independent of the stimulus contrast.

To understand theoretically the relationship between fractional paradoxical response and sub-circuit stability, we considered a 5-unit network (Fig. 3**c**, top right), with two PV populations, a perturbed one (red) and an unperturbed one (green). For each fraction of perturbed cells, the connectivity of this network is chosen such that its response to perturbations is mathematically equivalent to the mean response to that partial perturbation in the large-scale system under the HFA (see Sec. 5.9). As predicted by Eq. 1, whenever the response of the perturbed PV population in the 5-unit model system becomes negative (paradoxical), the sub-system composed of all of the non-perturbed populations loses stability; and, as in Fig. 2, in both the HFA and the fully nonlinear system, the outlier eigenvalues (those with largest real part) of the Jacobian of the sub-circuit of non-perturbed cells will be very close to those in the 5D system (Fig. 3**d**, top). Thus, when the mean response of the perturbed PV population in the HFA becomes negative, the sub-system without those perturbed neurons will become unstable (see Figs. 3**d**, S3; and Methods Sect. 5.9).

This understanding links stability of the non-perturbed circuit to the fractional paradoxical effect: whenever the system exhibits a fractional paradoxical effect, the unperturbed neurons will form a stable circuit for small numbers of perturbed cells, and will lose stability only after a critical fraction of cells are stimulated. Note that, at high contrast, there are networks for which the sub-system loses stability but for which the mean perturbed population does not change sign (Fig. 3**d**, bottom, upper right quadrant). The link between perturbation and stability is not bi-directional; the system can lose stability without changing the sign of the determinant of the Jacobian, and thus without evoking a paradoxical response (Eq. 1; see Miller and Palmigiano, 2020 for a clarification). In sum, although instability does not imply a paradoxical response, observation that a set of perturbed cells shows a mean paradoxical response tells us that the circuit without the perturbed cells is unstable.

### Theoretical foundation of VIP and SOM mediated disinhibition

VIP and SOM interneurons engage in competitive dynamics that regulate pyramidal cell activity via direct inhibition and disinhibition (Tremblay et al., 2016). We find theoretically that, when VIP projects only to SOM (as is at least approximately true in mouse V1, Campagnola et al., 2022; Karnani et al., 2016; Pfeffer et al., 2013, and as we observe in the connectivity matrix we inferred from data, Fig. 1), the response of E, PV and SOM to a change in the SOM input is negatively proportional to the response to a change in the input to VIP, with the same proportionality constant for all three cell types. VIP response shows the same proportionality, plus an additional direct VIP response to VIP drive. Thus, simultaneous drive to SOM and to VIP can cancel in their effects on E, PV, and SOM cells, causing only an increase in VIP activity. While drive to SOM or to VIP causes changes of opposite sign, it is theoretically possible for either one to have the positive sign, *i.e*. to evoke disinhibition. This depends on which is dominant, SOM inhibition of E cells or of PV cells, since the latter disinhibits E cells. This choice corresponds to a difference observed between layers in mouse V1. In layer 2/3, disinhibition is mediated by VIP: When VIP activity increases, it inhibits SOM, releasing E cells from its inhibition (Pi et al., 2013) and increasing E activity (Attinger et al., 2017; Fu et al., 2014; Karnani et al., 2016; Keller et al., 2020a; Mossing et al., 2021; Pi et al., 2013; Williams and Holtmaat, 2019). In layer 4, in contrast, SOM interneurons evoke disinhibition, because SOM projects preferentially to PV instead of E in this layer (Xu et al., 2013). Finally, although we can mathematically show this negative proportionality of responses only for the mean responses of each cell type, we find in simulations that this proportionality of response also holds true at the single cell level.

To establish these results, we start with the analysis of the 4-unit model, for a generic connectivity that satisfies the condition that VIP projects only to SOM but is otherwise arbitrary. We find mathematically (see Methods section 3.4) that the response of any given cell type to a VIP perturbation is negatively proportional to the response to a SOM perturbation, with a common proportionality constant across the four cell types (Fig. 4**a**, see also Eq. S15). In the case of VIP, there is an additional response component determined by its own gain. More concretely, if 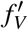 is the gain of VIP at a particular steady-state configuration (that is, the slope of its input/output function *f_V_*, which determines its firing rate at the steady state) and *ω_SV_* is the synaptic weight from VIP to SOM, then

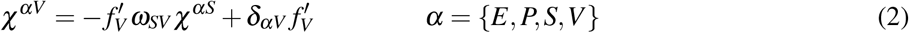

where *δ_αV_* is the Kronecker delta (*δ_αV_* = 1, *α* = *V*; = 0, otherwise).

**Figure 4.**
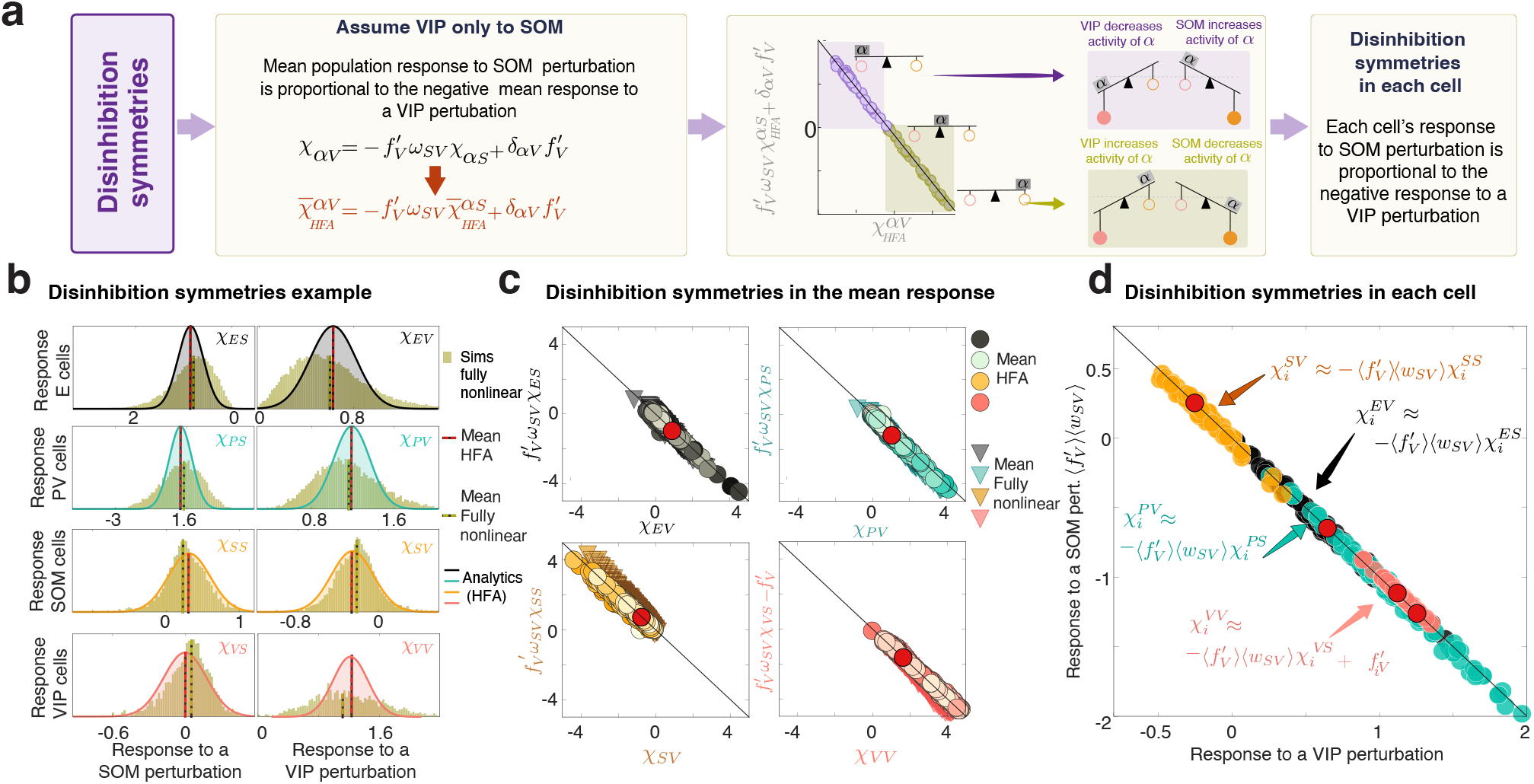
Disinhibition symmetries. **a)** Left: A parameter-independent relation holds true between the response of pyramidal cells (E), PV interneurons (P), SOM interneurons (S), and VIP interneurons (V) to perturbation of SOM and VIP under the assumption of VIP projecting only to SOM. Middle: The response of a cell type *α* to VIP perturbation, vs. the response to a SOM perturbation multiplied by the coefficient in Eq. (2), are negatively proportional. Two regimes can be identified: One in which VIP disinhibits while SOM inhibits the cell type *α* (lower right, green) and another one in which the opposite is true (upper right, purple). **b)** Distribution of responses to full-population SOM (left) and VIP (right) perturbations, for maximum stimulus contrast. Green histograms are the result of the simulation of a fully nonlinear large-scale system, while colored histograms and corresponding lines are the analytical results obtained using the HFA. Note the similarity of mean response in the fully nonlinear model (black dashed lines) and the HFA (red dashed lines). **c)** Opposite and proportional mean responses in the fully nonlinear system (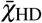, triangles) and under the HFA (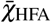, circles) to perturbations of SOM and VIP, for E (top left), PV (top right), SOM (bottom left) and VIP (bottom right), for the best 360 model fits. Because these data-compatible connectivities have only small values of connections weights from VIP to E, PV and SOM (Fig. 1**e**), they all show this symmetry between responses to SOM and VIP perturbations. These best-fit models all display VIP-mediated disinhibition and SOM-mediated inhibition of E and PV cells (compare E and PV responses to green quadrant in **a**). **d)** Disinhibition symmetries at the single neuron level. Responses of randomly chosen single cells (for the best model fit) of type E (black), PV (turquoise) and SOM (orange) and VIP (pink) cells to a perturbation of the entire VIP population (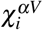), vs. the responses of those same cells to a perturbation of the full SOM population 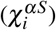 multiplied by the factor: 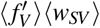 where the average is over the population (and for VIP cells, adjusted for self-excitation by VIP stimulation). The response symmetries hold at the single cell level in the fully nonlinear system.

We refer to these equalities as *Disinhibition Symmetries* (Fig. 4**a-c**). Because the mean response of a cell-type population to the perturbation of all neurons in another cell-type population is (under the HFA) given by the response of the 4-unit circuit (see Methods section 5.11), these symmetries also apply to the mean of the distributions in the large scale system under the HFA. Figure 4**b** shows the response distributions of an example model from Figure 1**d**, to perturbations of the entire SOM and VIP populations. The distributions obtained in simulations of the fully non-linear system (green histogram) are in good agreement with the analytic results under the HFA, and exhibit the disinhibition symmetries. Mean responses across the 360 models that best fit the data all accord well with the disinhibition symmetries (Fig. 4**c**), both in models under the HFA (colored circles) and the fully nonlinear networks (triangles), as expected since the data-compatible models all exhibit only weak connections from VIP to cell types other than SOM (Fig. 1**e**). All of these models predict the VIP-mediated disinhibition observed in L2/3.

Finally, we show that, surprisingly, the mathematical understanding provided by the disinhibition symmetries at the mean level, also apply at the single cell level. Figure 4**d** shows, for a single fully-nonlinear network example, that the response of each cell to a perturbation of the full SOM population is negatively proportional to the response of that cell to a perturbation of the full VIP population, with a proportionality factor approximated as the average connectivity from SOM to VIP (〈*w_SV_*〉) and the average gain of VIP cells 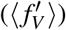. These responses are perfectly anti-correlated (see also Fig. S4 for log-normally distributed weights and connectivities in which VIP projects to all cell types).

The disinhibition symmetries formalize a clear intuition: Because VIP neurons only project to SOM neurons, a weak perturbation to VIP will only affect the rest of the circuit through SOM, relaying that perturbation with an opposite sign. The opposing, proportional effects on E, PV and SOM of small VIP versus SOM perturbations, with the same proportionality constant for all, are strong predictions resulting from our analysis.

### VIP-SOM cancellation underlies behavioral modulations of activity

We next asked, if behavioral modulations of activity act as an extra input –a perturbation– to the local circuit, what are the characteristics of those inputs? We addressed this inverse problem – asking what is the distribution of inputs targeting each cell-type population that would lead to observed changes in activity – to study the mechanisms underlying a known behavioral perturbation: locomotion. We examined the difference between each cell’s activity in the locomotion and the stationary condition in the experimental data (Fig. 5**a**). We found, consistent with previous findings (Fu et al., 2014; Pakan et al., 2016), that in the absence of visual stimulation the mean change in activity is only positive for VIP cells (Fig. 5**a** and **c**); that of E cells is not. This fact leads to a puzzle: why does the large change in VIP activity not cause an increase in E activity via VIP-mediated disinhibition as widely reported in layer 2/3, even in the absence of visual stimulation, (Mossing et al., 2021)?

**Figure 5.**
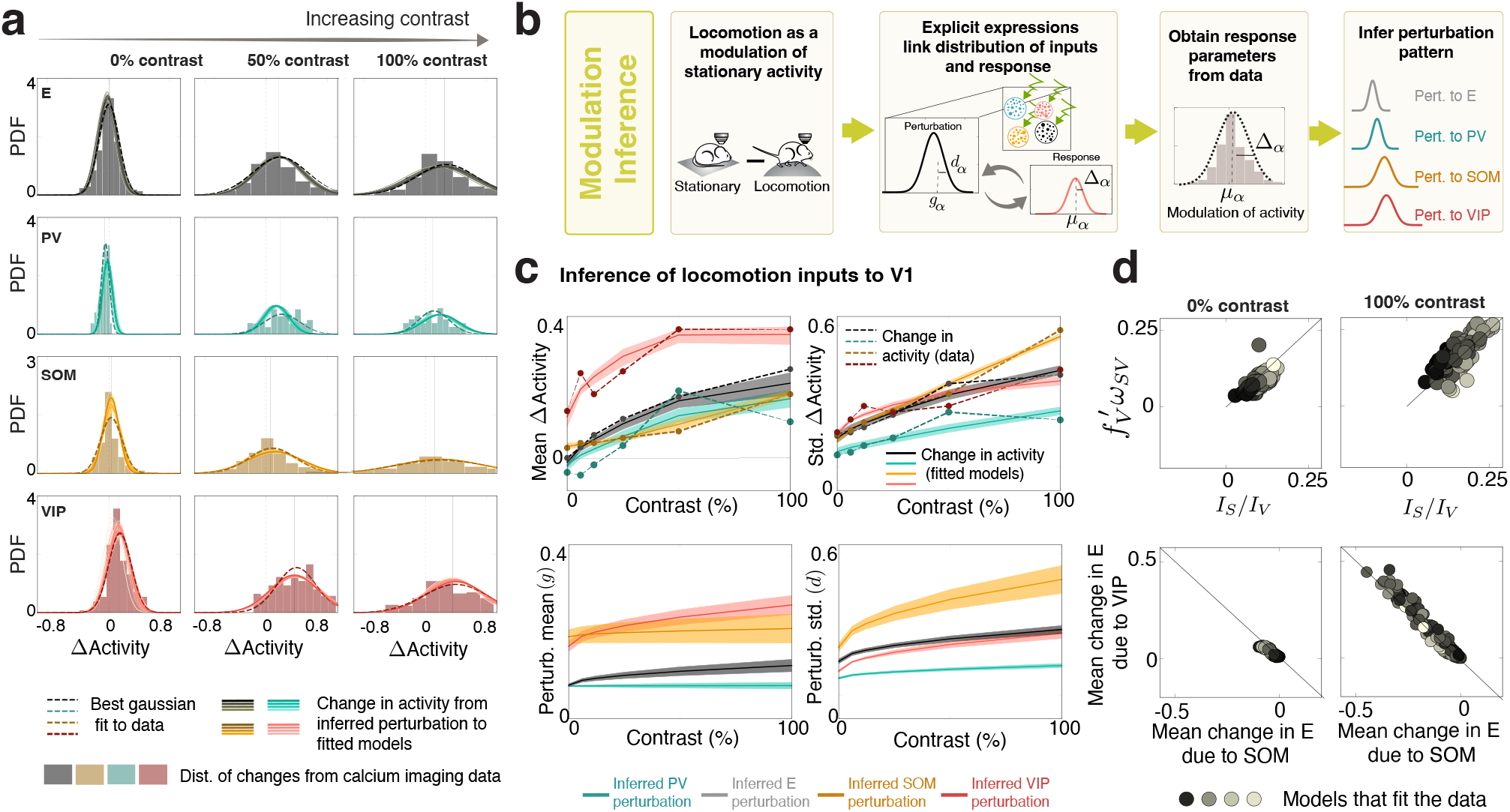
Inferring locomotory-related inputs reveals VIP-SOM cancellation. **a)** Distribution of Δ activity (difference between each cell’s activity in the locomotion and the stationary condition) for each cell type and different values of the stimulus contrast. Dashed lines indicate best Gaussian fits. Solid lines are fits from the explicit expressions (see Eqs. S126 and S129) **b)** Scheme of how to infer the cell-type specific perturbations (Gaussians of mean *g_α_* and standard deviation *d_α_* for *α* = E,PV,SOM,VIP) that give rise to the distributions of Δ activity (with mean *μ_α_* and standard deviation Δ_*α*_). **c)** Left two panels: Mean and standard deviation of Δ activity as a function of contrast. Dashed lines are the data (E in black, PV in blue, SOM in orange and VIP in dark red). Full lines are the mean over all model fits. Shaded areas indicate the 95 percentile of the distribution of those changes in activity over the model fits, obtained as described in **b**, for the family of models that fit the stationary data. Right two panels: The mean (g) and the standard deviation (d) of the inferred perturbations. The shaded area indicates the 5th through 95th percentile of the distribution of the values of the mean and std of the inputs over different models. **d)** The ratio of the mean inferred inputs to SOM and to VIP *I_S_/I_V_* are proportional to the mean VIP gain 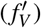 times the mean strength of the connection from VIP to SOM ((*ω_V_*), different for each model). This is the case even when the stimulus contrast is large and the mean change in E due to locomotion no longer vanishes. Bottom: Change in E activity due locomotory inputs to VIP Vs due to locomotory inputs to SOM.

To answer this question we expanded the framework described in previous sections to include heterogeneous perturbations (see Methods section 5.10). The obtained expressions, which depend on the stimulus contrast only implicitly via the population’s gain, allowed us to map the statistics (mean and variance) of a given perturbation, to the statistics (mean and variance) of the changes in the recurrent-network activity induced by that perturbation. By fitting these expressions to the changes in activity induced by locomotion (Fig. 5**c**, left panels), we could infer, for each model in the family of models that fit the stationary data (see Fig. 1), the distribution of inputs to each cell type population that would generate distributions of responses in the circuit identical to those produced by locomotion at each value of the stimulus contrast (Fig. 5**c**, right panels).

We found that, to account for the locomotion-induced activity changes, the inputs to VIP must have a large mean and variance, as discussed in Fu et al. (2014). Surprisingly, we found that inputs to SOM also had to be strong. This does not imply that SOM activity increases with locomotion. Instead, in the absence of visual stimulation, the effects of the direct locomotory input to SOM are largely canceled by inhibitory inputs coming from VIP. This is an example of the disinhibition symmetries described in the previous section. It is this cancellation at the level of the SOM output that leads to the zero mean change in pyramidal cell activity in the absence of visual stimulation.

We further found that this cancellation holds when the network receives visual inputs, across all contrasts. This is true even though the mean change in activity induced by locomotion and its variance increase with stimulus contrast for all cell-types (Fig. 5**c**, left panels). According to equation 2, for this cancellation to happen, the mean locomotory input to SOM *I_S_* and to VIP *I_V_* should fulfill (see Eq. S30).

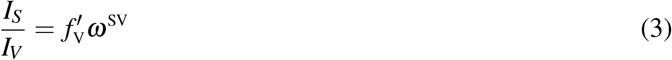

Plotting 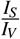 vs. 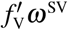 shows a scatter plot of this relation for all models that fit the data for all stimulus contrasts. We find that the inferred mean inputs cluster around the diagonal line, indicating that the inputs to SOM and to VIP largely cancel in their effects (Figure 5**d**, top). Directly examining the changes in mean E activity induced by the locomotory inputs to SOM and to VIP we find that these changes largely cancel, with only a few models showing weak disinhibition (Figure 5**d**, bottom). To infer the locomotory inputs, we assumed that the mean and standard deviation of the inputs varied as the square-root of stimulus contrast, but these results hold equally if instead they are assumed to vary linearly in, or to be independent of, the stimulus contrast (Fig. S5**b**).

If the effects of locomotory inputs to VIP and to SOM cancel, why does locomotion increase E firing rates in the presence of a visual stimulus? According to the inferred inputs, this is due to a small direct locomotory input to E cells, which increases with contrast (Fig. 5**c**, 3rd panel).

## Discussion

Experimental work has shown that multiple non-sensory signals are represented in areas previously thought to be highly specialized for sensory processing. Behavioral states such as thirst (Campagnola et al., 2022), hunger (Padamsey et al., 2022), or fear (Li et al., 2019) as well as multiple motor signals (Attinger et al., 2017; Bouvier et al., 2020; Dipoppa et al., 2018; Keller et al., 2012; Liska et al., 2022; Meyer et al., 2018; Miura and Scanziani, 2021; Musall et al., 2019; Niell and Stryker, 2010; Pakan et al., 2016; Stringer et al., 2019; Vélez-Fort et al., 2018), are represented in the sensory cortex of the mouse. Here we investigate the effect of two non-sensory signals, optogenetic perturbations and locomotion, in modulating cell-type-specific activity in the visual cortex of the mouse in the presence of different visual stimuli. We proposed a parallel between these modulations of activity, and developed a common theoretical framework to study responses to such cell-type-specific, full or partial perturbations in large-scale heterogeneous circuits. This framework is composed of three stages i) inference of heterogeneous, cell-type-specific recurrent model circuits from distributions of sensory responses; ii) characterization of responses to perturbations of the model circuits; and iii) inference of cell-type-specific inputs that can account for the distributions of activity changes induced by locomotion or other behavioral states.

We leveraged the mathematical tractability of large-scale recurrent network models to find a family of circuit models that match the cell-type-specific distributions of visual responses observed across stimuli of varying contrasts. These models are specified by the statistics (means and variance) of the connectivity between each pair of cell types and of the stimulus-specific inputs to each cell type. We found that, provided the network is constrained to be inhibition-stabilized (i.e., has strong, potentially destabilizing recurrent excitation, Ozeki et al., 2009; Tsodyks et al., 1997, as observed across mouse cortex, Sanzeni et al., 2020), this method robustly recovers a specific connectivity structure found in mouse V1: few or no connections of VIP cells to cell types other than SOM, or of SOM cells to themselves (Campagnola et al., 2022; Karnani et al., 2016; Pfeffer et al., 2013) (see Figs. 1 and S1).

We found that these models, despite spanning a broad distribution of parameters, respond similarly to perturbations, and that cell-type-specific responses reveal mechanisms of circuit stabilization. Specifically, we established that, if stimulation of all of, or a subpopulation of, a cell type evokes a “paradoxical” response – meaning that the stimulated cells show a negative mean steady state response to positive stimulation – then the circuit without the stimulated cells is unstable. In our data-fit models, PV cells, and not any other cell type, are circuit stabilizers and show a paradoxical response, consistent with other theoretical results (Bos et al., 2020) and with experimental findings that suppression of PV but not SOM cells leads to epileptiform activity (Veit et al., 2017). Furthermore, we predict a “fractional paradoxical effect” for partial perturbations of PV cells (or any cell type showing a paradoxical response to full-population perturbation): with increasing proportion of the population stimulated, the fraction of the population responding negatively changes non-monotonically. The fraction of negative responses decreases as a function of the fraction of stimulated cells until shortly before the stimulated population is big enough that the network is unstable without it, meaning that suppression of that population leads to epileptiform activity. This prediction can be directly tested by simultaneously monitoring (via calcium imaging) and perturbing (via holographic optogenetic stimulation) an increasing number of PV cells.

In layer 2/3 of the visual cortex of the mouse, SOM and VIP interneurons engage in competitive dynamics that underlie multiple forms of contextual modulations of activity (Keller et al., 2020a,b; Mossing et al., 2021; Veit et al., 2017). We showed that, because VIP projects almost exclusively to SOM cells (Campagnola et al., 2022; Karnani et al., 2016; Pfeffer et al., 2013), the effect of a VIP perturbation on the mean activity of other cell types is proportional to, but opposite in sign to, their mean response to SOM perturbation, with the same proportionality constant for all cell types (VIP cells show the same proportional response, plus an additional response to their own stimulation). This means that the effect of simultaneous inputs to both SOM and VIP can largely cancel for all cell types except VIP interneurons. Furthermore, while our theory established this cancellation at the level of the mean responses of each cell type, we found in simulations that this cancellation reliably characterized responses at the single-cell level. Remarkably, this cancellation is the mechanism that we found underlies locomotory modulations of activity.

We used our perturbative framework to infer the distribution of locomotion-induced inputs to the four cell types from their distributions of locomotion-induced response changes. We found that, to account for these response distributions of all four cell types, such a cancellation of VIP and SOM influences must be taking place. This cancellation accounts for the large increase in VIP firing rates while producing little change in other types. As a result, locomotory increases in excitatory-cell firing rates are inferred to be primarily due to direct locomotory drive to excitatory cells, which is consistent with the finding that acetylcholine, which is released in V1 in response to locomotion (Fu et al., 2014), increases the gain of thalamocortical inputs (Gil et al., 1997; Hsieh et al., 2000) (further discussed below). This contrasts with the hypothesis that locomotion-driven increases in E-cell visual responses are due to disinhibition caused by activation of VIP cells (Fu et al., 2014).

Note that experimental findings that activation of VIP cells increased excitatory firing rates, while suppressing VIP activity prevented the effects of locomotion (Fu et al., 2014), do not contradict our prediction: either manipulation imbalances the VIP-SOM competition, activating or suppressing the disinhibitory circuit and thus altering excitatory cell activity, yet those effects may not be driving the normal effects of locomotion.

Our findings are in line with experimental and theoretical work suggesting that locomotory inputs in V1 influence all of the cell types. These locomotion-related signals have multiple sources, each with different projection patterns to the different cell types. Cholinergic inputs from the basal forebrain, which have been shown to at least partially underlie the modulations of activity induced by locomotion (Fu et al., 2014; Lee et al., 2014, but see Polack et al., 2013, target many or all interneuronal types (Arroyo et al., 2014; Colangelo et al., 2019; Fanselow et al., 2008; Kawaguchi, 1997; Kuchibhotla et al., 2017; but see Yaeger et al., 2019). Noradrenergic input from the locus coeruleus, which targets multiple cell types (Kawaguchi and Shindou, 1998), also is implicated in locomotory response changes (Polack et al., 2013). Locomotory inputs can potentially also arrive to V1 from motor cortex, which sends direct projections to V1, mostly strongly targeting excitatory and PV cells (Leinweber et al., 2017). Finally, as locomotory inputs are already present in the lateral geniculate nucleus (Busse, 2020; Erisken et al., 2014), and locomotion-induced acetylcholine release increases the gain of thalamocortical synapses (Gil et al., 1997; Hsieh et al., 2000), we expect that visual drive to L4, and thus to L2/3, carries mixed locomotory and visual signals. In particular, an increase in gain of thalamocortical synapses could induce a multiplicative scaling of visual responses, consistent with our finding that the locomotory input to E cells should be increasing with stimulus contrast (note that we found that locomotory inputs targeting SOM and VIP can be independent of the visual stimulus, see Fig. S5).

The family of models we obtained describe the responses to small visual stimuli, averaged over stimulus orientations. This allowed us to ignore the distributions of neurons across space and preferred orientation and the dependencies of connectivity on differences in these features (*e.g*., Cossell et al., 2015; Ko et al., 2011; Rossi et al., 2020). Future work will be needed to extend the framework proposed here to embrace space, orientation, or other features on which connectivity may depend. It is in principle possible that larger visual stimuli, which strongly engage SOM cells (Adesnik et al., 2012), would lead SOM to play a role in stabilizing the circuit. However, optogenetic suppression of SOM cells in the presence of a large visual stimulus do not show signatures of instability (Veit et al., 2017), and we expect that the predictions proposed here are robust to changes in stimulus configuration.

Finally, the models developed here could be applied to study the differences in the effects of optogenetic and behavioral modulation across different cortical areas. Auditory cortex, for example, is an area that shares multiple organizing principles with V1, such as a VIP → SOM → E disinhibitory circuit (Bigelow et al., 2019), inhibitory stabilization (Kato et al., 2017; Moore et al., 2018) and SOM engagement in suppression by a stimulus surround (Lakunina et al., 2020). Nevertheless, the effect of locomotion in A1 is to suppress excitatory cells, rather than to enhance their firing as in V1 (Zhou et al., 2014), and this effect is not mediated by VIP (Bigelow et al., 2019). It is conceivable that, instead of a different mechanism underlying behavioral modulations of activity across sensory areas, differences in the degree of cancellation between SOM and VIP projections to the pyramidal cell population could lead to the observed differences. More generally, our work provides a general framework to study both behavioral and optogenetic modulations of activity and how those are shaped by the responses of sensory stimuli.

## Acknowledgements

A.P. would like to acknowledge the support of the Swartz Foundation Fellowship for Theory in Neuroscience 2019-4 and 2020-6. K.D.M., H.A., A.P., and D.P.M. would like to acknowledge funding from NIH 5U19NS107613. K.D.M. and A.P. also acknowledge funding from NIH U01-NS108683 and R01-EY029999, from NSF NeuroNex 1707398, and from Gatsby Foundation GAT3708. FF was supported by RIKEN Center for Brain Science, Brain/MINDS from AMED under Grant No. JP20dm020700, and JSPS KAKENHI Grant No. JP18H05432. D.P.M. was supported by an NSF Graduate Research Fellowship. H.A. is a New York Stem Cell Foundation-Robertson Investigator. H.A. and D.P.M. acknowledge the funding of NEI grant R01EY023756-01. This work was supported by the New York stem cell foundation. All authors would like to thank L. Abbott, B. Doiron, G. Handy, A.L. Kumar, and L. Mazzucato for useful feedback on earlier versions of this manuscript.

## Author Contributions

A.P. F.F. and K.M. conceived the study. A.P. and N.K. designed the low-dimensional fit approach. A.P. designed the high-dimensional fit approach and performed the numerical simulations and the analytical calculations with recommendations of F.F. and supervision from K.M.. D.M. recorded and analyzed all the experimental data under the supervision of H.A.. A.P., F.F and K.M. wrote the paper. All authors discussed the results and contributed to the final stage of this manuscript.

## 1 Methods summary

We develop a two-stage program for the prediction of responses to perturbations of large-scale circuits with multiple inhibitory types (Fig. 6). In a *first stage*, we build large-scale models that account for the entire distribution of cell-type-specific responses observed in the data. For that, we start by fitting a 4-unit dynamical system (see Sec. 3.1) to the mean responses observed experimentally in the four cell types (excitatory(E), PV, SOM, VIP) in layer 2/3 of the mouse visual cortex, in response to stimulation by a small (5 degree diameter) visual stimulus of varying contrast (for data details see Sec. 2), while the animal is in a stationary condition. These fits are obtained by finding parameters that are a solution to a non-negative least squares (NNLS) problem (Renner et al., 2020) (see Sec. 3.2). These simple models allow for a mathematical understanding of the response to perturbations to different cell types, and to make parameter-independent predictions (see Secs. 3.3, 3.4 and Fig. 6**c**). Then, we use these 4-unit model fits as a parameter seed to build a family of large-scale models (Fig. 6**d-e**, see Sec. 4) that captures the full distributios of activity for each cell type and stimulus contrast. To find the parameters of these models we leverage their mathematical tractability (see Sec. 4.4). First, as the distribution of activity has a tractable analytical form (Roxin et al., 2011) that depends on the mean and variance of the input currents to each cell-type population, we can obtain that mean and variance for each cell-type and stimulus condition by fitting that distribution to the data via maximum likelihood (Fig. 6**f**, left). This gives us the target mean and variance of the input currents for each cell type that the model needs to be able to self-consistently create. Given some model parameters (Kadmon and Sompolinsky, 2015) (*e.g*., means and variances of connection strengths and external inputs), the means and variances of the input currents can be computed self-consistently through Mean-Field (MF) theory (Fig. 6**f**, right). Now, to find the parameters that make the model capture the data, we explicitly write the Kullback-Leibler divergence between the activity distributions that fits the data and the distributions that the model creates. By sampling parameters from distributions created from the 4-unit fits, and by choosing a suitable threshold on this distance (0.5) (Didelot et al., 2011), we find large-scale models whose distributions of activity across cell types and stimulus contrasts reproduce those observed experimentally (Fig. 6**f**, middle). These fits make predictions about the mean and variance of the connection strengths between neurons of any two given cell types, and of the external inputs each cell-type receives as a function of the visual stimulus.

**Figure 6.**
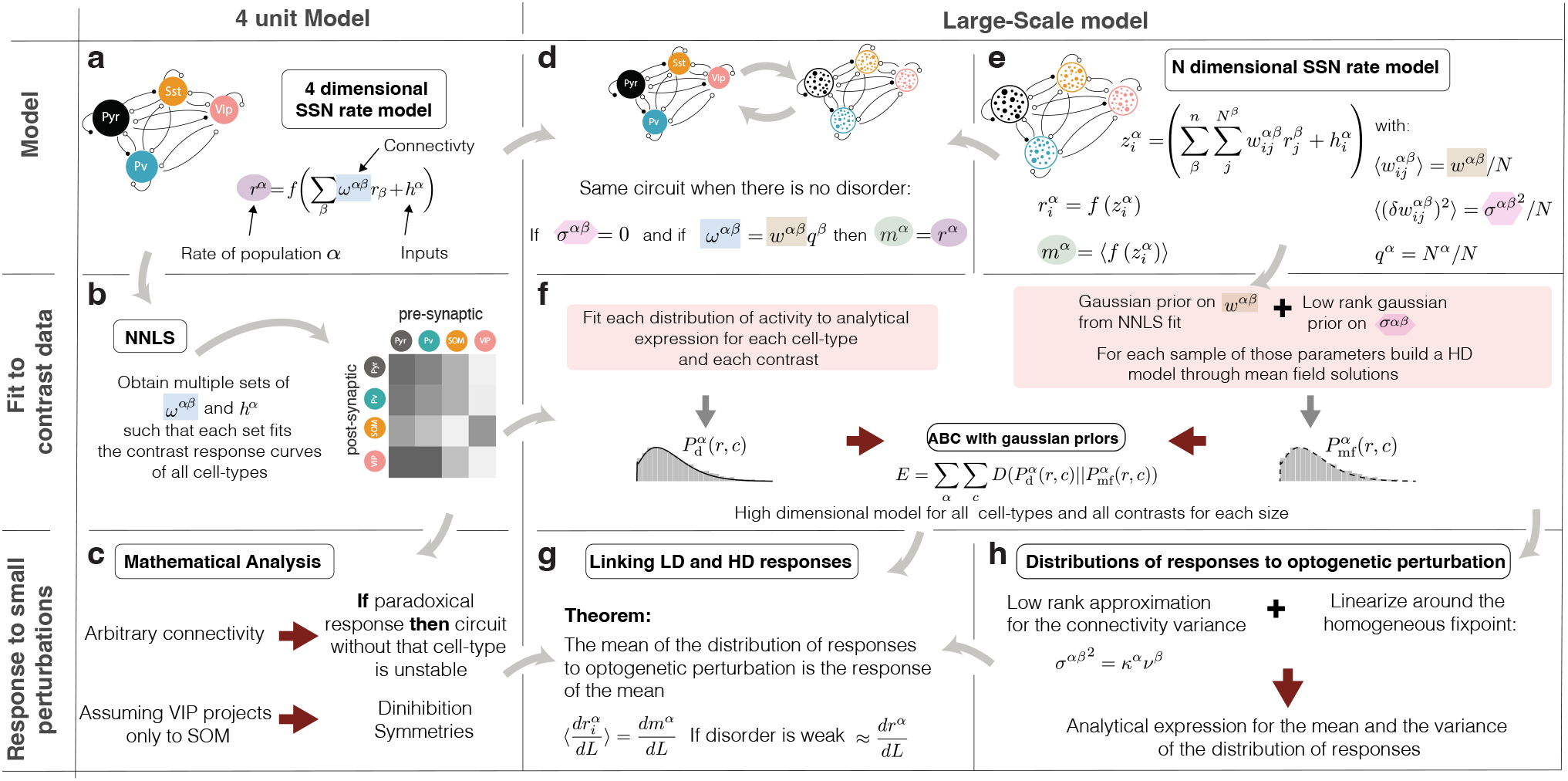
Outline of this paper. **a-b)** 4-unit circuit): circuit describing the mean activity of E, PV, SOM, and VIP cells when presented with stimuli of different contrasts. By using non-negative least squares (NNLS) we find the parameters to fit the contrast response of all four cell types. **c)** We find relations between stability and responses to optogenetic perturbations and, assuming that VIP only projects to SOM and SOM does not project to itself, find hidden structure in the response matrix. These findings are applied to the models that fit the data. **d-e)** Large-scale model: There are now many neurons of each cell type. When there is no disorder - all the connections of a given type (as defined by presynaptic and postsynaptic cell types) have the same strength, and all cells of a given type receive the same external input, the large-scale circuit describes the same dynamics as the 4-unit circuit described in (a), provided that the parameters are chosen appropriately. Inclusion of disorder changes the mean activity. **f)** We use Approximate Bayesian Computation, also known as ABC, to fit the high-dimensional system. Firstly, the models we use have analytical expressions for their distributions of activity. We fit such an expression to the measured distribution of activity of each cell type for each stimulus condition. Secondly, we build large-scale models with parameters sampled from a distributions with priors on the means obtained from the NNLS analysis. For each model, we use mean field theory to compute the resulting activity distributions. By minimizing the Kullback-Leibler divergence (Cover and Thomas, 2005) between these two sets of activity distributions (the one obtained from the data and the one obtained from the model family), we find the models that best approximate the activity distributions of all cell types at all contrasts with a single parameter set. **g-h)** Analytical expressions for the distribution of responses to optogenetic perturbations are available for linear systems. Through an approximation, we linearize the high-dimensional system and compute the entire distribution of responses to an arbitrary pattern of optogenetic stimulation.

In a second stage, we build on random matrix theory results (Ahmadian et al., 2015) (see Sec. 5.1 and 5.3) to analytically compute, under the homogeneous fixed point approximation (HFA, i.e. that all neurons in each cell type’s population have the same gain, see Fig. 6**h** and Sec. 5.2), the distribution of neuronal responses to patterned perturbations. We show that under this approximation, the mean response to an excitatory perturbation to all cells in a cell-type population in the large-scale system has a mean given by the response of the system without disorder to that same perturbation (see Sec. 5.5). Furthermore, this response is itself mathematically identical to the response of an equivalent 4-unit system (see Sec. 5.4 and Fig. 6**g**). This means that the parameter-independent predictions of the response to perturbations in the low dimensional case (4-unit model), carry over to the mean response to a full population perturbation in the large-scale system. Furthermore, under the HFA, partial perturbations lead to a bimodal distribution of responses given by a mixture of Gaussians (see Sec. 5.7), whose mean (see Sec. 5.5) and variance (see Sec. 5.6) can be computed exactly.

We rely on a Theorem by T. Tao (Tao, 2013) to link the stability of the network subsystems and the paradoxical response of a given cell-type. First we show that in the 4-unit system, if the sign of the response of a cell-type to its own perturbation is negative, that implies that the circuit without that cell-type is unstable. Under the HFA, what can be shown is that the bulk eigenvalue distribution lies in a ball centered at 1, and contains only stable eigenvalues (those with real part < 0); and that, by Tao’s theorem, there are the outlier eigenvalues are given by the eigenvalues of the 4-unit system (see Sec. 6), which may be unstable. In other words, the outlier eigenvalues are the ones determining the stability of the system. Therefore, the insights about the relationship between stability and paradoxical responses obtained in the 4-unit system also carry over to the large-scale model under the HFA. Furthermore, as the response of the large circuit to partial perturbations is given by a mixture of Gaussians, and the mean of each Gaussian composing the mixture can be described by the response of a 5 dimensional circuit (four non-perturbed cell-type populations and a perturbed population, see Sect. 5.9), the link between response to perturbations and stability can be also studied in the case of practical perturbations.

## 2 Data Collection and Analysis

All the data presented here was collected by Daniel Mossing, refer to Mossing et al. (2021) for details about data collection. We analyzed the activity of 2664 pyramidal cells, 57 PV cells, 203 SOM cells and 169 VIP cells, whose receptive field was within 10 degrees of the stimulus center. The analyzed conditions were small drifting gratings (5 degree) for different contrast conditions (0%,6%,12%,25%,50%,100%). The cells were selected to have a significant response that differed from the baseline (pvalue<0.05). The activity shown in Figure 1 is the mean over trials and stimulus orientations. We did not select cells that were aligned to the orientation of the stimulus. While the mouse was free to run, the analysis focused on stationary data fundamentally, i.e. trials running speed was defined smaller than 1 cm/sec as in (Mossing et al., 2021) for all figures except in Figs. 5 and S5, where we took the difference between locomotion and non-locomotion conditions. In this case, the locomotion threshold was taken to be 5 cm/sec.

## 3 Low-dimensional circuit models (LD model or 4-unit model)

We consider a network of *n* = 4 units, each describing the activity *r^α^* of a particular cell type’s population *α*, with *α* = {E, PV, SOM, VIP} in layer 2/3 of the visual cortex of the mouse.

### 3.1 Definition

The network integrates input currents *z^α^* in the following way

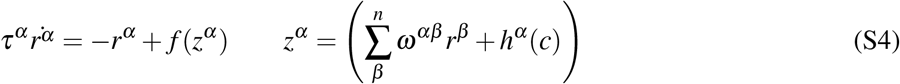

where *τ^α^* is the relaxation time scale, *ω^αβ^* is the connectivity matrix, and 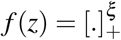 is the activation function. We use *ξ* = 2 unless otherwise specified. The inputs *h^α^* (*c*) are composed of a baseline input *h_b_*, a sensory-related input *h_s_*(*c*). This last input is chosen to be proportional to the contrast *c*, for which *h_s_*(*c*) = *h_c_c*, with *h_c_* a contrast independent variable to be fitted:

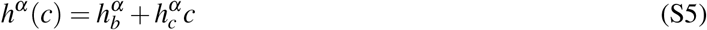

### 3.2 Fitting of mean activity

To simultaneously fit the mean rates of all four cell-types at all contrast values (six in total *c* = {0 6 12 25 50 100}), we consider the steady-state equations corresponding to (S4). Since the recorded firing rates are positive and non-vanishing, the inverse is well defined 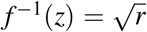 and the nonlinear steady-state equation corresponding to (S4) becomes a linear equation with respect to the connectivity parameters:

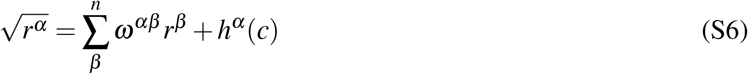

As stimulus-related inputs are linear functions of the stimulus contrast multiplied by different weights to E and to PV, *h^α^* = *h^E^, h^PV^*, 0, 0. Eq. (S6) represents a system of linear equations *Ax* = *y*, where *x* is an unknown vector containing the flattened connectivity matrix entries *ω^αβ^* and the input constants 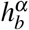 and 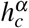. The entries of the matrix *A* and the vector *y* are the functions of the recorded firing rates at six contrast values. For each cell type, the matrix *A* has 6 rows corresponding to each of the six contrast values. The number of columns of the matrix *A* is equal to the number of unknown connectivity and input constants for the cell type *α*. This case in which the number of equations (rows of *A*) and the number of parameters (all chosen weights and inputs) are equal, the system *Ax* = *y* can be solved exactly. To be concrete, for each cell type *α*, we will have:

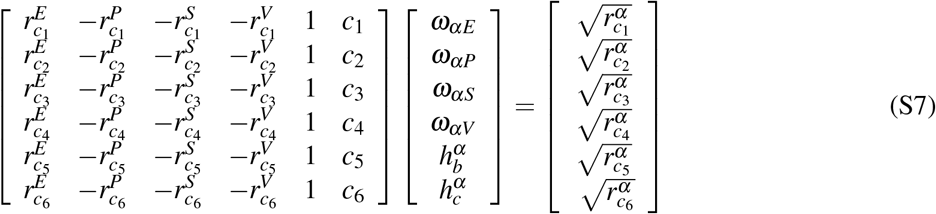

In the work presented here we expanded the above matrix A to solve all 24 equations (6 × number of cell types). Values of parameters that approximately solve the Eq. (S6) can be found by computing the non-negative least squares (NNLS) (Chen and Plemmons, 2009) solution. The NNLS solution of Eq. (S6) constructed from mean firing rates, gives *one* set of connectivity and input parameters *x*. To obtain distributions of connectivity and input parameters instead, we created surrogate contrast responses sets by sampling from a multivariate Gaussian distribution with mean 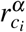 and standard error of the mean 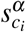. For each input configuration, we sampled 2500000 seeds to create these surrogate contrast response curves. For each sample contrast response *k*, NNLS gave one connectivity and input parameter set. Using each parameter set and the steady-state equations in (S6) we computed the fit 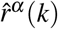 of the *k*th sample contrast response. Keeping the stable solutions (negative eigenvalues, all time constants were chosen to be equal to 1), the likelihood of that parameter set *k*

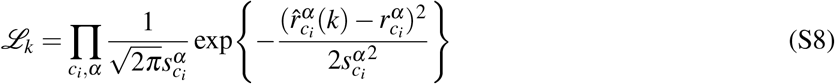

defined a hierarchy for the contrast response samples. From the family of LD models that fit the data, we only considered those that were ISN. We did not enforce any connectivity weights to be zero. Some of our models had also absent connections from SOM to VIP, we disregarded those. Models shown in Figure 1**a** are the top 200 of the 700 models that later were used as prior *seeds*.

### 3.3 Linear response and paradoxical effects

The linear response matrix is defined as the steady state change in rate of a population *α* given by a change in the input current *h* to population *β*

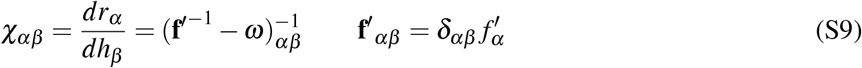

where 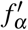 is the gain of population *α* at the considered steady state, **f**′ is the 4-dimensional diagonal matrix with elements 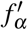, *δ_αβ_* is a Kroenecker delta which is 1 only if *α* = *β* and zero otherwise, and *I* is the 4-dimensional identify matrix. Defining the diagonal matrix of time constants *T_αβ_* = *δ_αβ_ τ_α_*, Eq. (S9) can be written as a function of the Jacobian *J* = *T*^−1^(−*I* + **f**′*ω*), and in particular of its negative −*J* = *T* ^−1^(*I* − **f**′*ω*), as

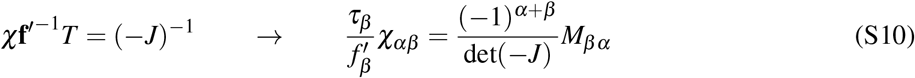

where we have used the general formula for the inverse of a matrix, and *M_βα_* is the corresponding minor of −*J*, that is, the determinant of −*J* with its *β^th^* row and *α^th^* column deleted. In particular, the diagonal entries of *χ* are

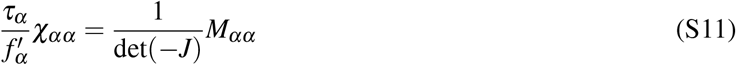

We let *J_α_* be the Jacobian of the sub-circuit without cell type *α*. Then *M_αα_* = det(−*J_α_*). Thus we find that:

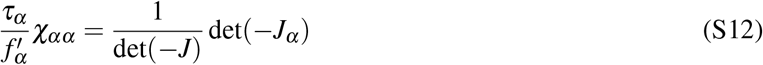

We assume the fixed point is stable, which means all of the eigenvalues of *J* have negative real part, so all the eigenvalues of −*J* have positive real part and det(−*J*) > 0. Similarly, if the subcircuit without *α* is stable, det(−*J_α_*) > 0, while if det(−*J_α_*) < 0, that subcircuit is unstable. Thus, if cell type *α* has a paradoxical response, meaning that *χ_αα_* < 0, then det(−*J_α_*) < 0, meaning that at the fixed point, the subcircuit without *α* is unstable.

#### 3.3.1 EI networks

Evaluating Eq. (S12) in the EI case we obtain the result from (Tsodyks et al., 1997)

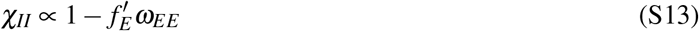

which makes the parameter independent prediction that when recurrent excitation strong (i.e. its 1D Jacobian is positive), the response of inhibition is paradoxical, *χ_II_* < 0.

#### 3.3.2 SOM paradoxical response implies unstable E-PV sub-circuit (VIP projects only to SOM)

In the particular case in which VIP projects only to SOM, the Eq. S12 reduces to

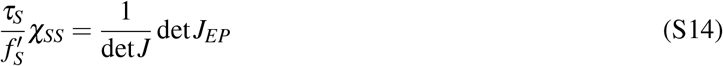

In EI networks, the network is stable if and only if the trace of the Jacobian is negative and the determinant of the Jacobian is positive. As in this networks the E time constant can always be chosen sufficiently large to satisfy the conditions on the trace, the stability will be given by the determinant alone. Therefore, measuring the paradoxical response of SOM translates in E-PV being unstable, and observing a non-paradoxical response of SOM means that E-PV is stable given a suitable time constant.

### 3.4 Disinhibition symmetries (VIP projects only to SOM)

The values *χ_αβ_* for the particular case in which the connections from the VIP population to the rest is exactly zero can be found to satisfy the following relations, *Disinhibition symmetries*.

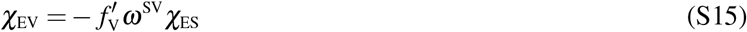

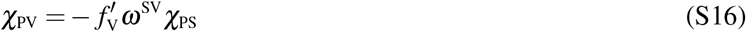

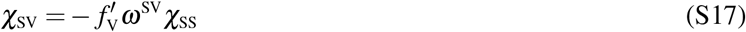

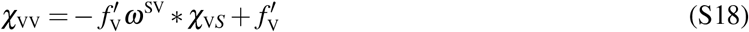

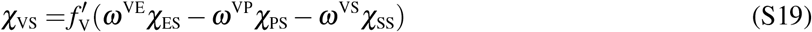

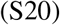

This can be easily seen by explicitly writing the response matrix as

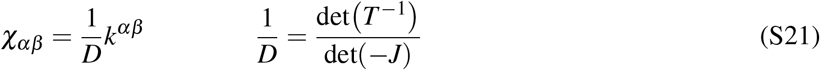

Where det(−*J*) is the determinant of the negative Jacobian of the full system, defined above Eq. (S10). Given that the eigenvalues of *J* have to be negative for linear stability, det(−*J*) is always positive, and the above relations can be instead written as a function of *k^αβ^* with

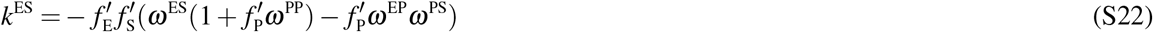

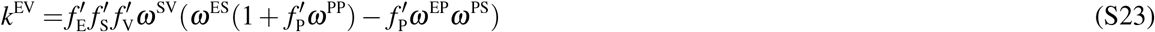

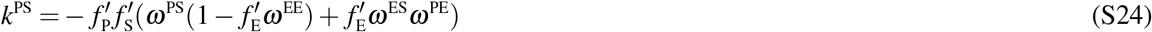

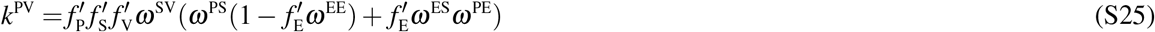

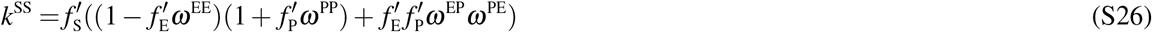

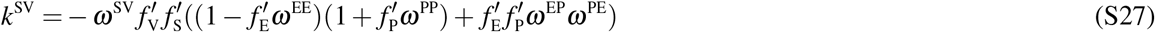

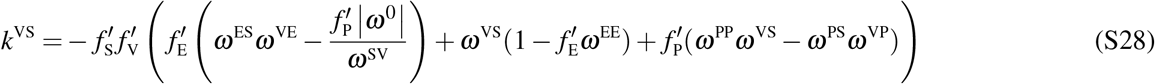

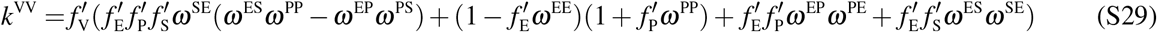

### 3.5 Disinhibition symmetries underlying behavioral modulations of activity

We find that, in the absence of visual stimulation, locomotory inputs target only SOM and VIP. In that case, for the change in E cell activity to vanish, its needed that the change in E activity due to the locomotory input to SOM (Δ*E_S_*) and the changes in E due to the locomotory inputs to VIP (Δ*E_V_*) cancel one another. Using the notation above and introducing the size of the locomotory inputs to S and V as *I_S_* and *I_V_* respectively, the locomotory inputs that would lead to such cancellation should respect:

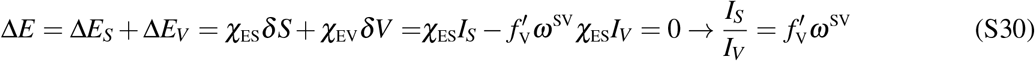

### 3.6 Equivalent class for low-dimensional circuits

To understand how the conclusions derived here would be modified by considering firing rates instead of deconvolved calcium imaging data we follow (Khan and Hofer, 2018), where it is reported that calcium activity 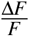 and firing rates can be related via a linear relationship. In general, given a power law input-output function 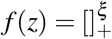, we can define a class of equivalent models by redefining activity together with weights and inputs

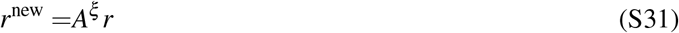

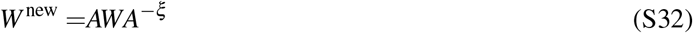

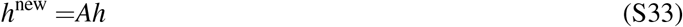

Where *A* is the diagonal transformation matrix from calcium activity *r* to firing rates *r*^new^. The Jacobian and the linear response matrix of this new system are related by:

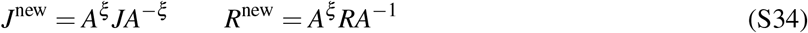

In particular given that the new and old Jacobian are related by a similarity transformation, this change of variables (or the equivalence class) will not change the stability. The the linear response can have re-scaled values but will preserve sign, and the *Disinhibition symmetries* equations will be re-scaled.

## 4 High-dimensional circuit models

### 4.1 Definition

In this section we describe the high-dimensional network models. The network has *n* = 4 populations with *N^α^* neurons in each population *α* = {*E, PV, S,V*}. We denote the fraction of neurons in each population by *q^α^* = *N^α^ / N*, where *N* is the total amount of neurons in the network. We took this fraction to be *q* = [0.8, 0.1, 0.05, 0.05] as is approximately in biology (Tremblay et al., 2016). The steady-state activity 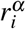 of the unit *i* in the population *α* is given by:

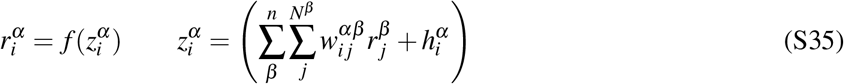

Whereby 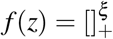 with *ξ* = 2 represents the transfer function of the neuronal populations. The connectivity elements 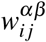 are Gaussian distributed with mean and variance defined by:

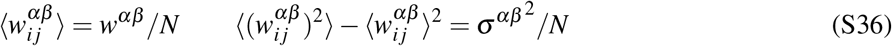

The inputs to each unit 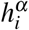 are also Gaussian distributed with mean 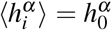 and variance 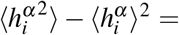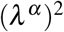. The steady state Eq. (S35) can be re written as a function of the input to each cell :

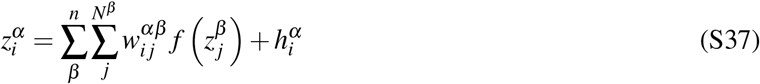

### 4.2 Set-up and mean field equations

In order to compute the mean and variance of the activity in each population self-consistently, we follow the approach in Kadmon and Sompolinsky (2015). The input 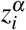 to a cell can be described as fluctuations around a mean: 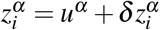. We define:

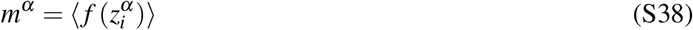

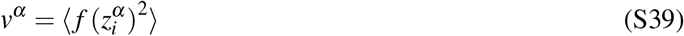

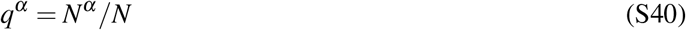

By taking the mean and the variance of Eq. (S37) and incorporating the definitions above, we re-obtain the self-consistent equations for the mean and the variance of 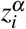, given by *u^α^* and Δ^*α*^

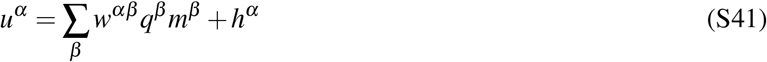

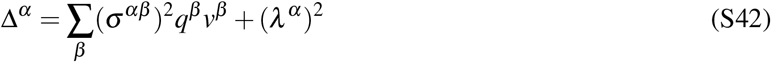

where

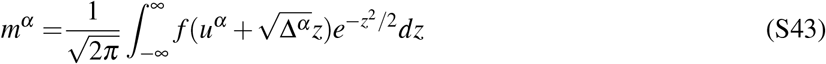

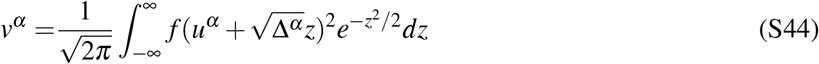

We observe that if there is no disorder, Eqs. (S41) and (S43) reduce to the model from Eq. (S4) with *ω^αβ^* = *w^αβ^ q^β^* and *m^α^* = *r^α^*.

### 4.3 Mean field perturbation

If *L* is a *homogeneous optogenetic* perturbation to the entire population *α*, the change in response of each cell is given by

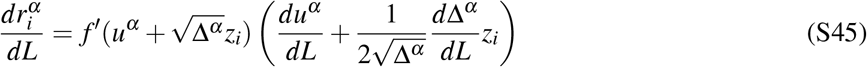

Taking the average and using Eq. (S43), we find that the mean of the response distribution to laser perturbation is given by the change in the mean activity of the population:

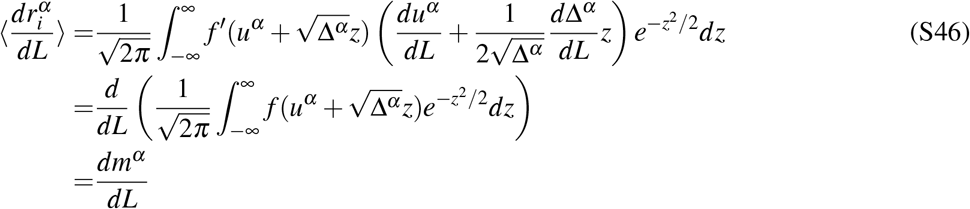

This equation relates how the mean of the distribution of responses to perturbation relates to the response of the mean activity in the fully nonlinear system.

### 4.4 Data Fitting

We emphasize that, due to the nonlinear transfer function, heterogeneity in the values of the synaptic connectivity will change the mean activity compared to the system without heterogeneity. Consequently, it is not sufficient to use the parameters found for the LD model as the mean values of the heterogeneous connectivity and input distributions; the means and variances of the connections and the inputs have to be found simultaneously for the large scale model to fit the data.

To fit the system defined by Eqs. (S41, S43), we used that the distribution of activity of a population *α* with the transfer function of the form 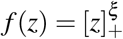 can be written (when assuming that inputs are Gaussian distributed) as a function of the mean total input *u^α^* and its variance Δ^*α*^ (Roxin et al., 2011):

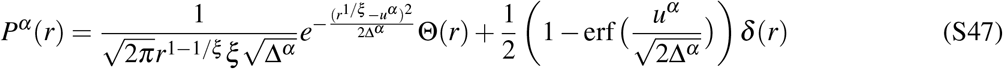

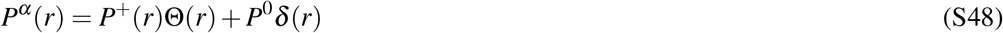

Here, Θ and *δ* denote the Heaviside and delta functions, respectively.

To find the parameters that approximate the distribution of the experimentally recorded activity we use Eq. S47 with *ξ* = 2 and proceed as follows (Fig. 1**a**, see also Fig. S1): For each cell type *α* and each contrast *c*, we fit the analytical distribution of rates from Eq. (S47) to the distribution of experimentally recorded activity. We denote the fit distribution by 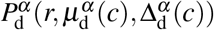. The fitted distribution 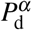 provides us with an estimate of the mean, 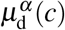, and variance, 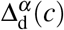, of the total input to each cell type and each contrast *c*. We assume that the mean external input to the population *α* has the form 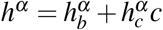, and has a standard deviation *λ*^α^ which is independent of contrast. All the cell-type receive baseline inputs but only E and PV have contrast dependent inputs. The model has in the end 34 parameters (16 mean weights and 8 low-rank variances of the weights, 4 mean baseline inputs, 2 mean weights for the stimulus-related inputs to E and PV, and 4 input variances, independent of contrast).

To find which parameters *w^αβ^*, *σ^αβ^*, *h^α^* and *λ^α^* best fit the data we proceed as follows: we do Approximate Bayesian Computation, also known as ABC search (Sisson et al., 2018), from prior distributions for the mean and variance of the weights and inputs to this network to build multiple instances of 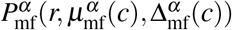.

The priors for *w^αβ^* and 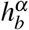 and 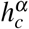 were Gaussian distributions with mean given by the parameters of the LD fits and a 5% std. The priors for *σ^αβ^* and *λ^α^* were chosen to be Gaussian distributions with mean 0.5 and variance 0.3. The only dependence on contrast is through the inputs mean, the variance of the inputs is independent of contrast. This was enforced to capture the dynamical spreading out of the activity with contrast (see Figure 1) and to not enforce it with an input that already is broader with contrast. Nevertheless, it is possible that the inputs received by layer 2/3 cells from layer 4 are already broader at higher contrast.

We define an error that depends uniquely on 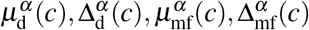. Specifically, we define the total error as the squared norm of the matrix of the Kullback-Leibler divergences between these two distributions:

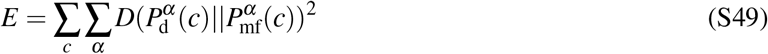

where D (dropping temporarily the dependence on the contrast for ease of notation) is given by:

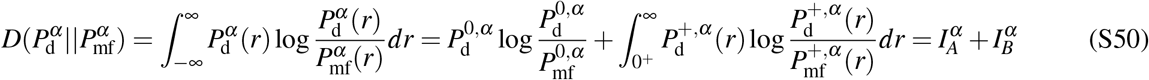

with

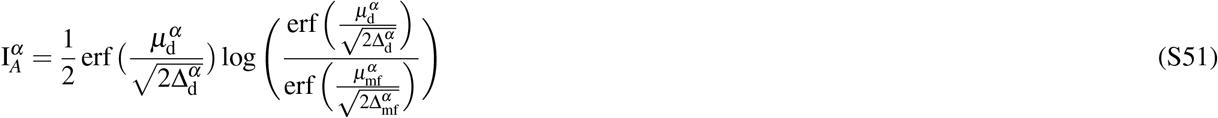

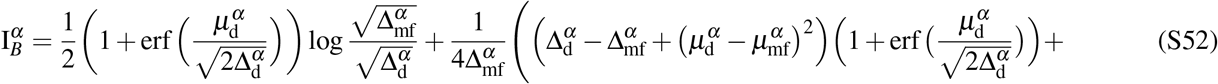

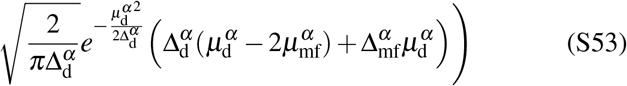

Instead of following the gradient to find an optimal solution we keep the solutions that have a sufficiently small error from the random sampling. Randomly sampling from these priors we obtained 500000 models whose total KL divergence was 0.7. From those, we either take the first 300 or those with KL divergence smaller than 0.5. This defines a family of high-dimensional models (Fig. 1) with skewed distributions that are in good agreement with the calcium activity, and capture not only the nonlinear dependence of the activity mean but also spreading out with increasing contrast.

### 4.5 Equivalent class for high-dimensional circuits

Analogously as shown for the LD case above, we are interested in understanding how to map the models derived here to those we could have derived had we had access to firing rates recordings instead of deconvolved calcium imaging data. We assume that calcium activity 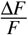 and firing rates can be related via a linear relationship, as shown in (Khan and Hofer, 2018). Let us say we want to change the mean activity of each cell type *α* by a factor *x^α^*

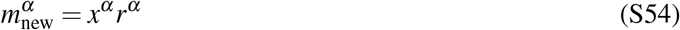

Then from Eqs. (S43, S41) we have

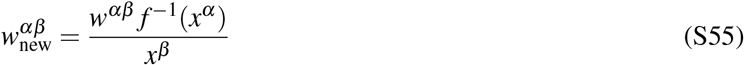

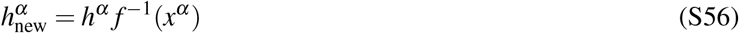

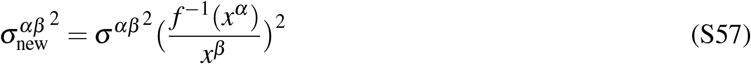

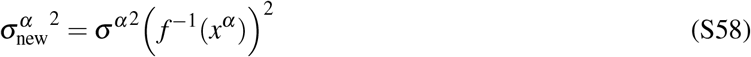

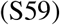

## 5 Analytical approach to linear response of disordered networks

### 5.1 Set up

We call the steady state solution of Eq. (S35) 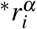 and the steady state input 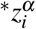. The time evolution of the response to a perturbation 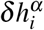, can be described by the dynamics of 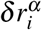:

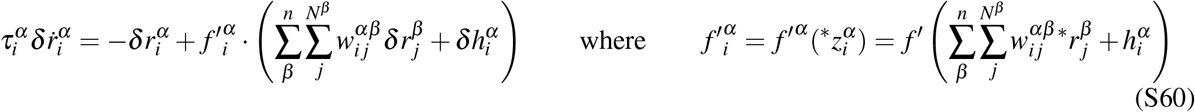

Switching from now onwards to matrix notation, we define

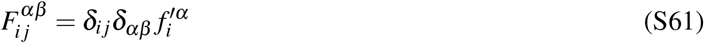

the diagonal matrix of derivatives, where *δ* is the Kronecker delta, and 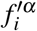 is the gain of neuron *i* in population *α*. The connectivity matrix *W* has elements 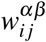. The steady state response to an arbitrary increase in the input given by *δh* will be:

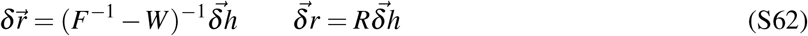

which defines the high-dimensional linear response matrix *R* = (*F*^−1^ − *W*)^−1^. In order to be able to compute the block-wise variance of the response matrix analytically, we need to constrain the cell-type specific variance to be low rank. The block-wise variance of *W* (defined in Eq. S36) is then written as (*σ^αβ^*)^2^/*N* = *ν^α^ κ^β^ /N*. We write *W* as the sum of a homogeneous component *W*_0_, with 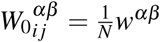 and a random component Π_*L*_ΨΠ_*R*_, where Ψ is a matrix with Gaussian distributed random numbers with zero mean and unit variance, and Π_*L*_ and Π_*R*_ are non-random diagonal matrices:

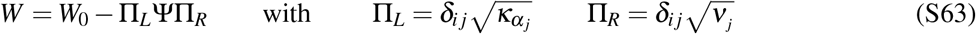

### 5.2 Homogeneous fixed point approximation (HFA)

The mathematical treatment we are going to outline later is only possible in linear system in which the disorder does not affect the gain of each neuron. That approximation, which we call the Homogeneous fixed point approximation or HFA, assumes that the linearized system can be written as :

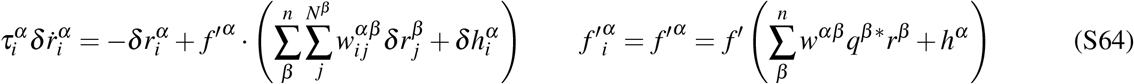

In practice, we solve the non-disordered system to compute *f′^α^* and look at the response matrix of that system. We observe that the HFA will approximate the response of the nonlinear circuit to small perturbations but wont approximate the activity itself.

### 5.3 General framework to compute the linear response in networks in the HFA

Using results from (Ahmadian et al., 2015) we find that in the special case of the HFA, where 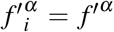 as described above, the mean linear response matrix over different instantiations of the disorder is the linear response of the non- disordered case:

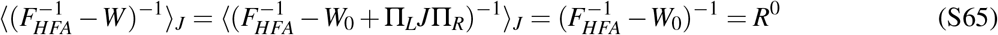

This fundamental relation links the mean of the distribution of responses to the response of the non-disordered system, its general in linear networks and works as a useful approximation in this case of study. Generally, in experiments, we have a perturbation pattern *δh* describing the proportion of stimulation each neuron receive, and a measuring vector *δb*, describing which are the neurons contributing (linearly) to the signal that we are going to be monitoring 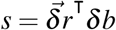. We compute the mean and variance of the signal *s* across different instantiations of the disorder. By defining:

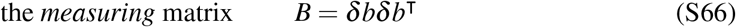

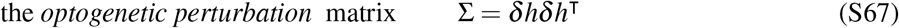

we can write the second moment of that measured signal *s* (Ahmadian et al., 2015):

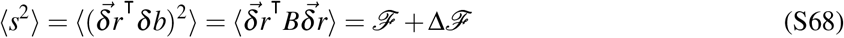

where

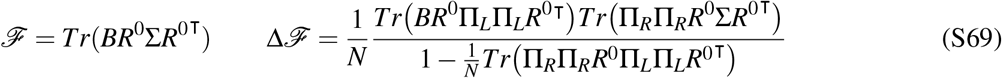

Where we used the definitions in Eq. (S63). We observe that in the absence of disorder, in which *W* = *W*_0_, 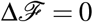 and the recorded signal is given only by *s* = *δbR^0^δh*^T^.

In the case in which we are interested in looking at single neuron statistics, we have *δb* = *e_i_* with *e_i_* = {0,…, 1, …, 0}

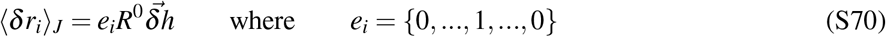

Eq. (S70) means that for each neuron, the distribution of linear responses over different instantiations of the connectivity has a mean given by the linear response in the absence of disorder (due to Eq. (S65)) and the variance Λ_*i*_ given by

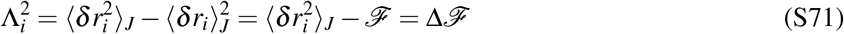

Equations (S65), (S68) and (S69) are general formulas for how to compute the mean and the variance of the linear response distributions as a function of the optogenetic perturbation Σ and the observation matrix *B*. In the following sections we will explicitly compute the mean response matrix *R*^0^ for both full and low rank connectivity and the variance in different optogenetic perturbation configurations.

### 5.4 Computation of the response matrix *R*^0^ **without disorder**

For computing *R*^0^ (given by Eq. (S65)) we write the block-structured matrix *W*_0_ as a function of the LD system connectivity *ω^αβ^* = *w^αβ^ q^β^*. We define the *N × n* matrices *V* and *U*, with columns given by *N* dimensional vectors

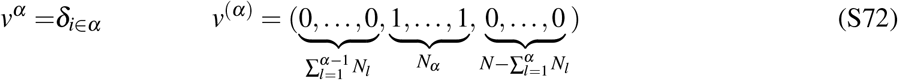

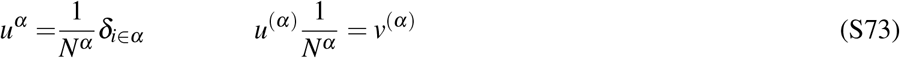

We can then write:

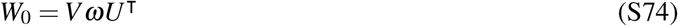

Where *w^αβ^* and *q^α^* were introduced in Eqs (S36) and (S40), respectively. To obtain *R*^0^ defined in Eq. (S65) we are going to exploit the fact that this is a low rank matrix. Depending on whether *ω* is also low rank or not, we will need to consider different strategies.

If *ω* is invertible, we can use the Woodbury lemma to find a succinct expression for *R*^0^:

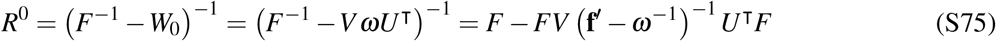

Introducing the notation *α_i_* as the population to which the neuron *i* belongs to, the entries of the response function can be written as

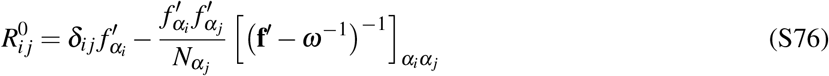

**f**′ was defined in Eq. (S9). We note that for this expression to be valid, *ω* needs to be invertible and in particular full rank. We also note that this expression is given by two terms: the first one, private to each neuron, is only non-zero if we are observing the same neuron that we are stimulating, while the second term, depends on which population the stimulated neuron belongs to and which population the observed neuron belongs to, but is independent on whether the perturbed neuron is the observed one.

We define 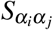, the sum of the linear response of a single neuron in population *α_i_* to a homogeneous input to the neurons in population *α_j_*

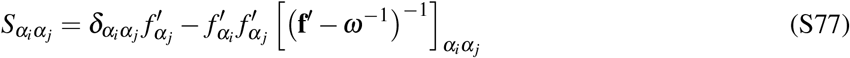

Substituting Eq. (S77) into (S76), we obtain an expression for the linear response which will be useful in later sections:

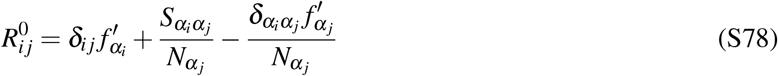

We point out that the Eq. (S77) is independent of *N*, and is finite in the limit of large *N*.

#### 5.4.1 Case of rank-one *ω*

In the case of rank-one *ω*, we cannot invert *ω* in Eq. (S75). Instead we write:

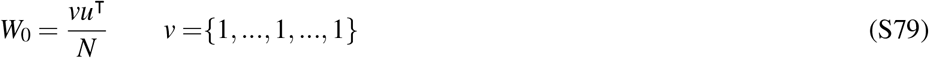

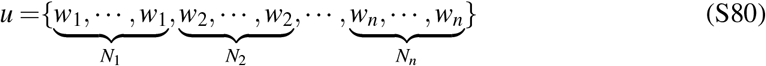

Using Sherman–Morrison formula we find that

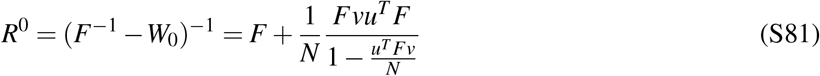

Where the denominator is always positive given that 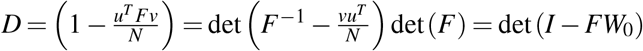. We obtain

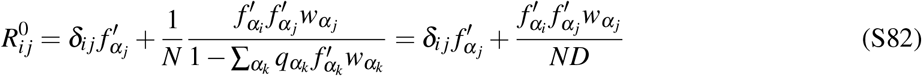

Again defining 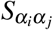 as the sum of the linear response of a single neuron in population *α_i_* to a homogeneous input to the neurons in population *α_j_*

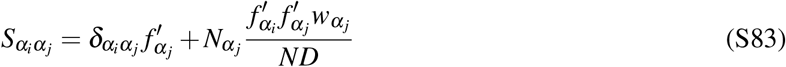

We note that the above expression is also finite in the large *N* limit. Using (S82) and (S83) we conclude that the linear response 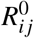 satisfies Eq. (S78) also in this case.

### 5.5 Response distribution to partial (homogeneous) perturbations: Mean term

In this section we consider fractional perturbations of neural populations, i.e. perturbations applied only to a subset of neurons in each population. We derive a formula for the sum of the linear response of a single neuron in a population *α_i_* to perturbations applied to fractions 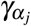 of neurons in populations *α_j_*. Within each perturbed population *α_j_*, we denote the set of perturbed neurons by 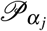. If we perturb 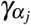 neurons in a population *α_j_* then we find that the response of the neurons in population *α_j_* that were stimulated have a mean response that depends on whether they were directly stimulated or not (see below).

If *ω* full rank (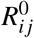 is given by (S76)), if we perturb *γ_η_* neurons in populations *η* the total perturbation vector is given by *δh* = {*δh*_1_, *δh*_2_, · · ·, *δh_n_*}, where 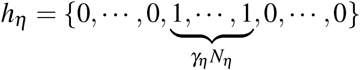. Then we find that the response of the neurons is given by

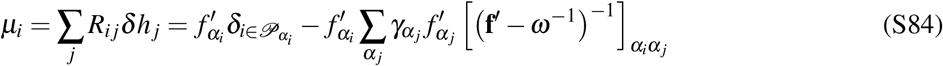

The expression in Eq. (S84) can be represented as a sum of the mean responses of directly perturbed and non perturbed neurons. The mean response of directly stimulated neurons is given by

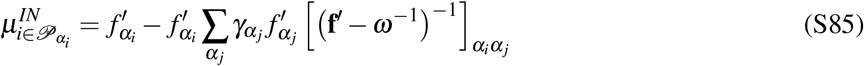

whereas the neurons in *α_j_* that were *not* stimulated and the neurons from other populations follow the equation:

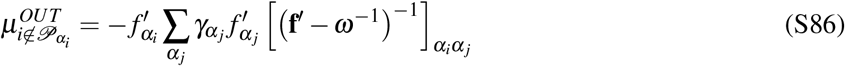

We note that these expressions critically depend on the sign of (**f**′ − *ω*^−1^)^−1^. To capture Eq. (S85) and Eq. (S86) as a single equation we define a matrix

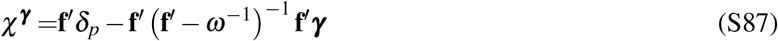

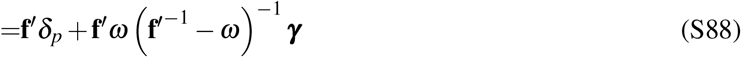

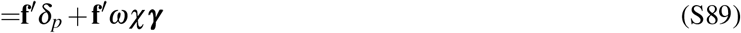

where *δ_p_* = 0 if we are describing the mean of the non-perturbed population and *δ_p_* = 1 otherwise.

In the case when we study the paradoxical response, meaning that we perturb and record activity in the same population we find that using 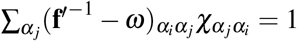 (matrix times its inverse is the identity) we have that 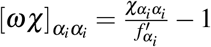. We rewrite (S84) as

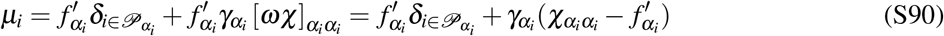

If the response is paradoxical in the LD system 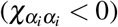, the response distribution of non stimulated neurons has a negative mean, and becomes even more negative if the fraction of perturbed increases. If the above term is positive for a small fraction of perturbed cells, it can become negative when the fraction of the perturbed cells increases. We denote the critical fraction of perturbed cells for which the response mean becomes negative by 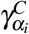 and obtain

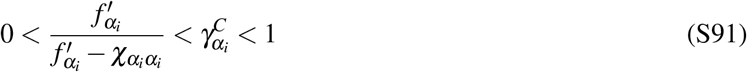

In Fig. 5, the *fractional paradoxical* effect occurs while the perturbed cells are not responding paradoxically. Nevertheless, before the mean of the distribution of perturbed cells changes sign, the distribution itself shifts left and therefore this critical fraction is different from the critical fraction for which the system is exhibiting a *fractional paradoxical* effect.

#### 5.5.1 Case of rank-one *ω*

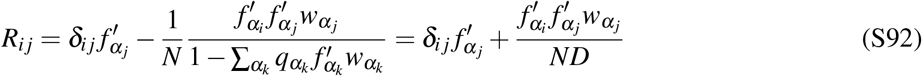

Identically as above, neurons that are directly stimulated will have a response given by

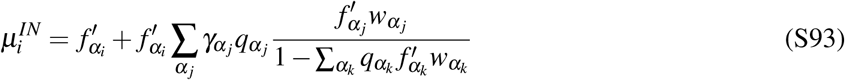

whereas the neurons in *α_j_* that were *not* stimulated and the neurons from other populations follow the equation:

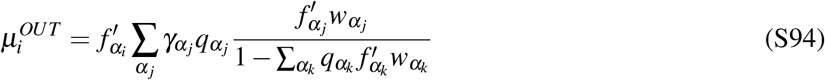

##### 5.5.1.1 Critical fraction

In the case in which we only have and *EI* circuit, and we stimulate only the inhibitory population, we can see that for inhibitory neurons in which *w_j_* is negative, the response of the neurons that were not stimulated is always paradoxical (meaning that Eq. (S94) is always negative), but the response of those neurons that were stimulated will only be paradoxical when 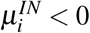

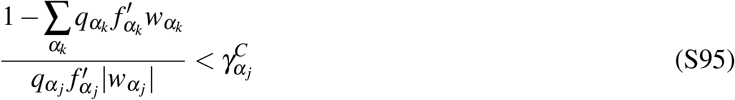

First let us consider the case in which we have a fixed amount of neurons but we have an increasing amount of populations *n*. Given that 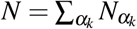 if we take 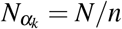 then 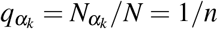. We find that the critical fraction in (S95) is now

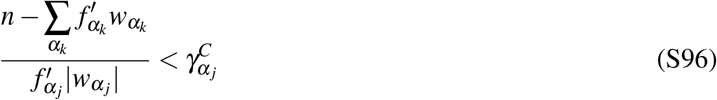

We find that given a fixed sum of *w_i_* and fixed N, the fraction of stimulated neurons *γ_k_* needs to increase linearly in *n* to have a paradoxical response.

##### 5.5.1.2 Comparison with Sadeh et al. (2017)

If now we would normalize the weights in Eq. (S79) as in (Sadeh et al., 2017), meaning that *w_j_/N* → *w_j_*/(*N/n*) (See Eq. 2 of their paper), the above equation would be

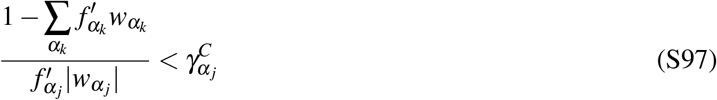

Taking 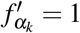 (linear network) we recover their result below Eq. 12 of their paper.

### 5.6 Response distribution to partial (homogeneous) perturbations: Variance term

From Eq. (S69) we know that the variance of the response is going to depend on the response of the system without disorder *R*^0^. The goal of this section is writing *R*^0^ in the form expressed in (S78). We will first find the general expression and then evaluate for particular cases: For that we write (S69) as

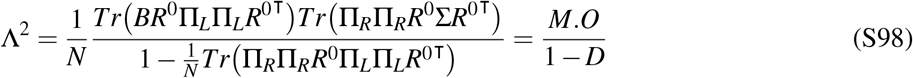

Where the *optogenetic targeting* matrix Σ = *δhδh*^r^. If we write *δh* = {*δh*_1_, *δh*_2_, · · ·, *δh_n_*}, where *δh_η_* is the perturbation vector for the population *η*, then for each *δh_η_* we can write 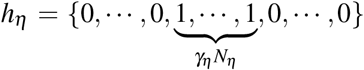, meaning that given *n* populations, there will be a vector with entries *γ_η_* that tells us which is the fraction of neurons of each population that we are stimulating. Each element of the *optogenetic targeting* matrix will then be:

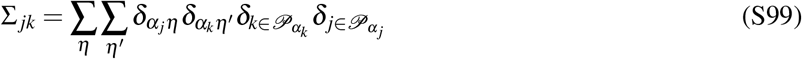

Observation: The *optogenetic targeting* matrix has entries in the off diagonal terms.

To compute the expression of the variance when we need to make three different terms defined above in Eq. (S98): The measuring term *M*, the optogenetic perturbation term *O* and the Denominator term *D*

#### 5.6.1 The measuring term *M*

Using the definitions in Eq.(S63), we write

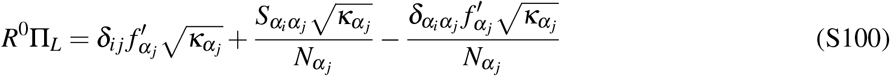

and

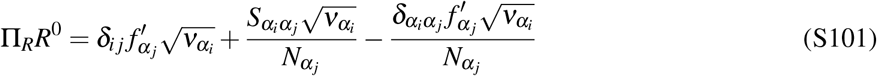

such that

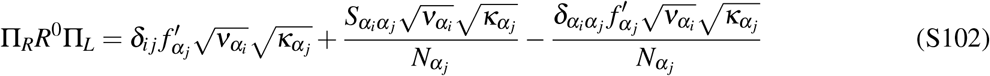

Therefore, the term we are interested in obtaining will be

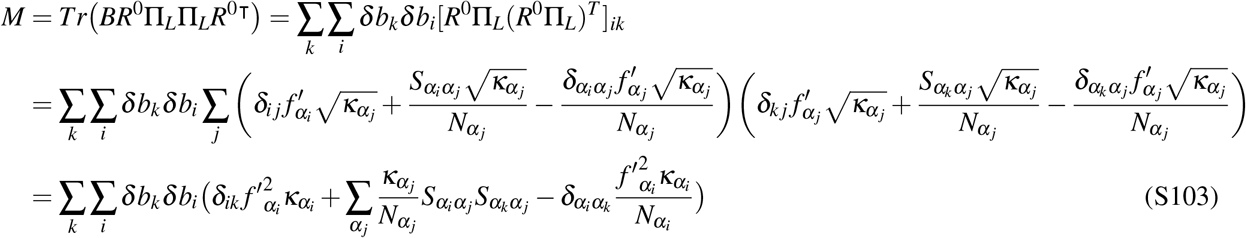

Which cannot be further reduced without specifying the specific shape of *δb*. In the articular case in which we are observing a single neuron, then *δb* = *e_l_* = {0, · · ·, 0, 1, 0, · · ·, 0} at some *l*. In that case *δb_i_* = *δ_il_* and

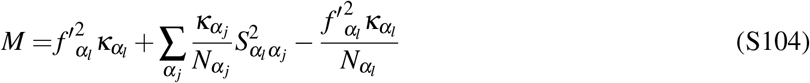

#### 5.6.2 The optogenetic perturbation term

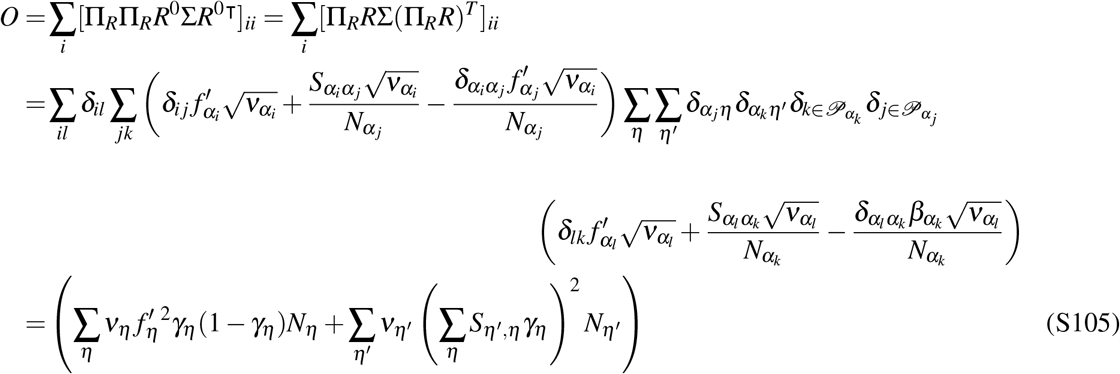

#### 5.6.3 The denominator

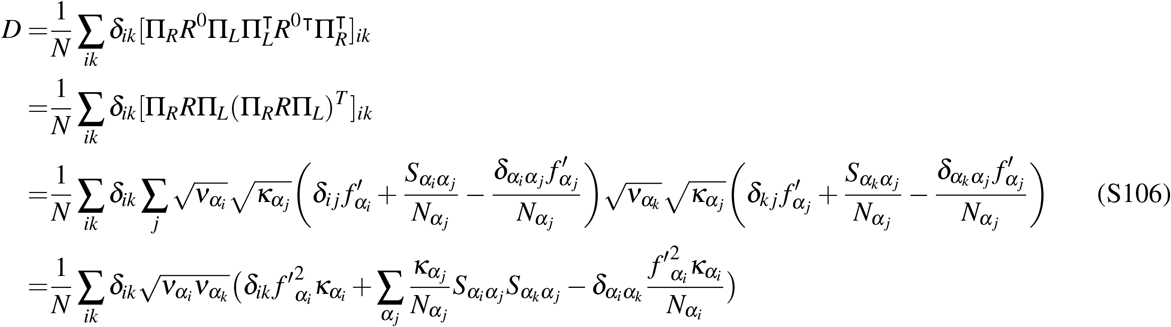

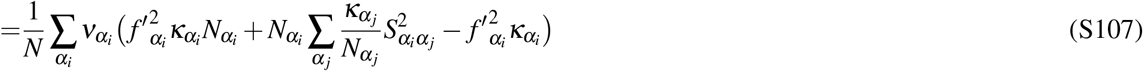

#### 5.6.4 The final expression

We write down here the final expression for the variance of the response of a single neuron in population *α_l_* while perturbing a fraction *γ_η_* of the population *η* (*q_η_* = *N_η_ / N*):

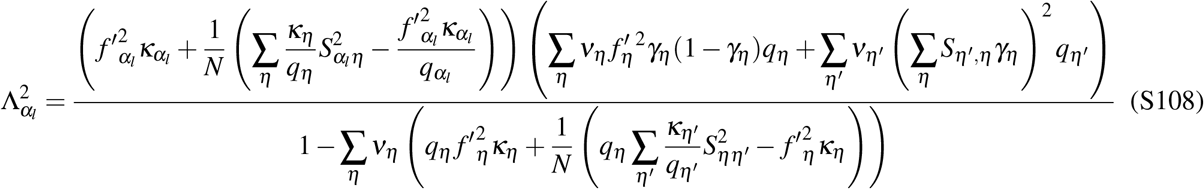

We observe that in the large N limit the expression reduces to:

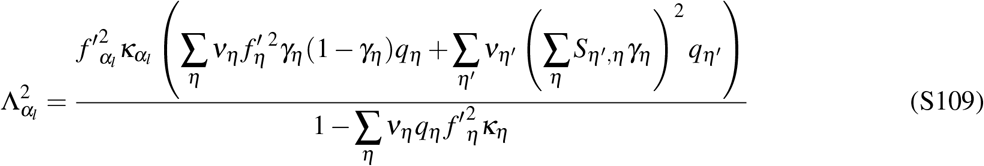

Which is independent of N iff *γ_η_* is a finite fraction of the population. In the case in which a finite amount of neurons *k* are stimulated, *γ_η_* = *k/N_η_* and the variance will vanish in the large N limt.

An interesting prediction is a nonlinear dependence of the variance of the populations with increasing fraction of stimulated neurons. The expression Eq. (S109) has a nonlinear term in the fraction of stimulated neurons in each population. When more than a single population is stimulated, there is also a term that nonlinearly mixes the fraction of interacting neurons. This results in non trivial dependences of the variance with the fraction of stimulated cells. Depending of the fraction of stimulated cells, the effect of increasing fraction of one-cell-type stimulation can be to narrow down the distributions or to broaden them. We name this a *second-order paradoxical effect*, which is different from the *fractional paradoxical* effect, which is an effect due to the mean changes.

##### 5.6.4.1 Simplification: Non-structured variance

In the particular case in which the degree of disorder on the connectivity does not depend on the pre- and the post-synaptic cell type, i.e. when *κ_α_* = *ν_α_* = *σ* we obtain a simpler expression for the variance of the populations:

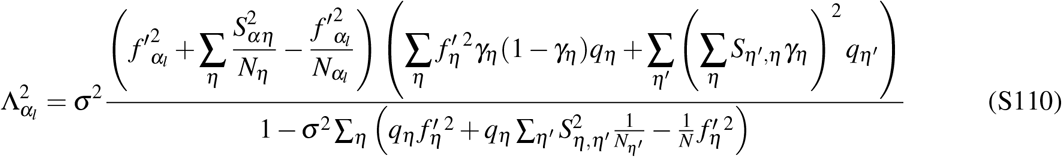

### 5.7 Response distribution to partial (homogeneous) perturbations: Full Distribution

In the case of partial stimulation, we will have a total distribution that is a mixture of Gaussians with means

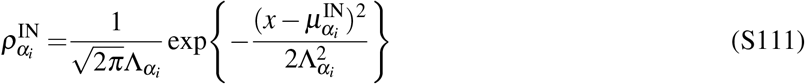

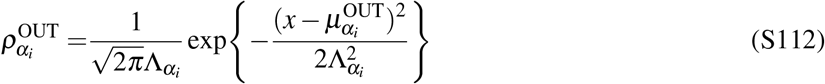

So the total distribution of responses is

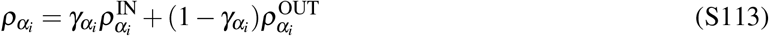

Where 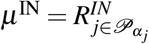 and 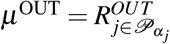 given by Eqs (S93, S94) for low rank *ω* or by (S85,S86) for invertible *ω* and a variance given by Eq. (S108)

### 5.8 Simple description of the fractional paradoxical effect

The fractional paradoxical effect can be intuitively understood in the system without disorder (the EI case with low-rank *ω* was studied by (Sadeh et al., 2017)). In this case, the distribution of responses will be bimodal, with two delta functions at the values given by Eq. (S90). The density then will be given by the limit of vanishing variance of Eq. (S113)

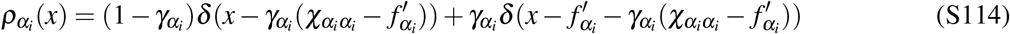

If the unit *α_i_* is paradoxical in the LD system, then 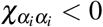. The left peak will always be negative, and for sufficiently small 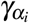 the peak of the perturbed cells will be positive. As computed in Eq. (S91), for values of 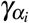 smaller than 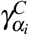, the mean of the perturbed population will remain positive. In this range, increasing the fraction of perturbed cells, will result in a decrease of the mass of negative responses 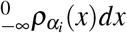 like (1 − *δ*). In the non-disordered case, as soon as 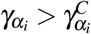, the mass of negative responses is unity. Given that when working with the homogeneous approximation, the response of the non- disordered system is the mean of the response of the disordered system, the intuitions here apply to the mean of the disordered case.

### 5.9 Fractional paradoxical effect and link to a 5D low-dimensional system

Here we show that the mean response of the perturbed fraction of the population can be mapped to the response of a system with 5 dimensions, in which the *α_i_* population, that here for simplicity we take to be PV, is split in a one perturbed and one non-perturbed populations. We know that mapping a high-dimensional non-disordered network to a LD system can be done by rescaling the weights according to the fraction of cells in that population. That manipulation will not change the activity of either cell type given that they receive the exact same input currents. The linear response of that system in consideration is, *χ*^5^ is given by

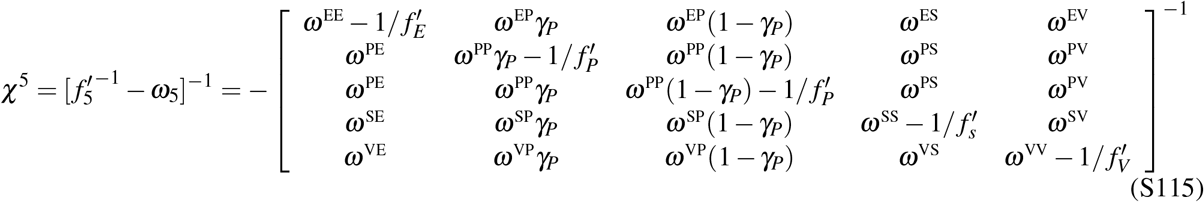

The term we are interested in is the PP term, given by:

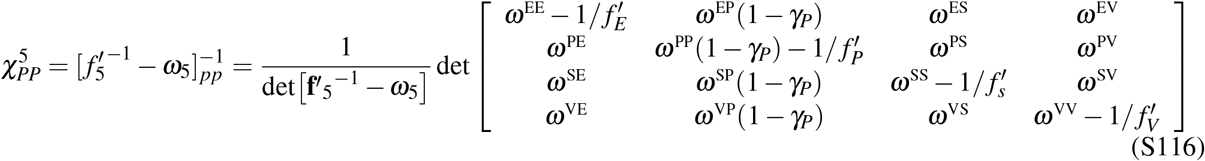

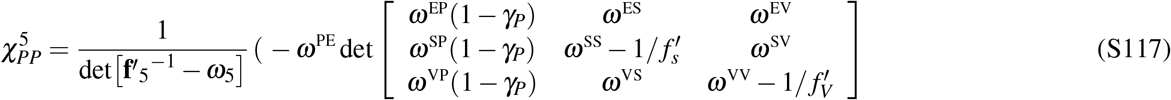

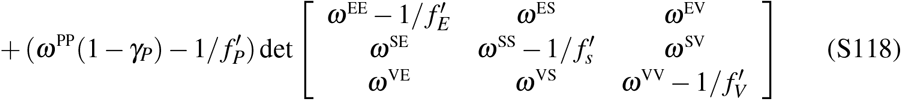

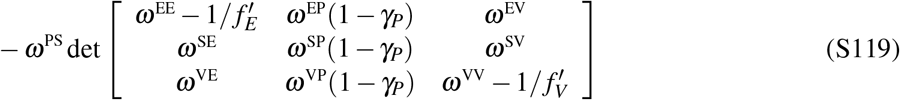

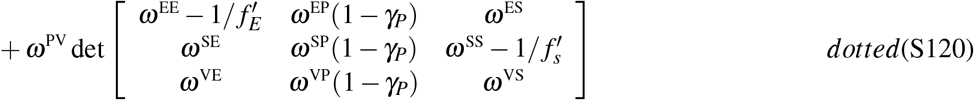

Each 3D determinant is *minus* the minor, M, of the original 4D matrix (**f**′^−1^ − *ω*). Using that

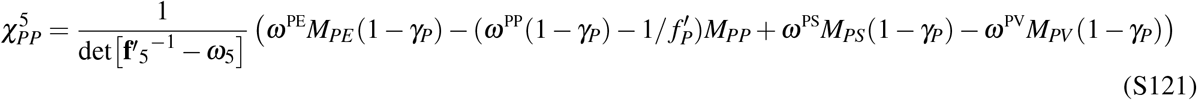

Where *M_αβ_* are the minors of the original 4D matrix (**f**′^−1^ − *ω*). Using that 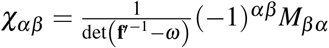, and that det 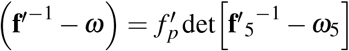

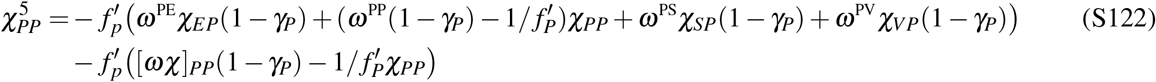

Using again the trick that 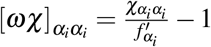

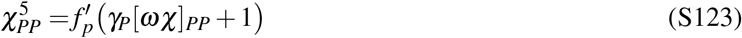

Given that the mean response of the perturbed population in a high-dimensional system, given by Eq. (S84) (and also Eq. S87) is 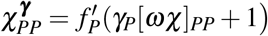, we obtain that

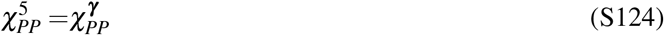

And as both determinants are positive because of linear stability, this two things have the same sign. This calculation, together with Eq. (1), tells us that whenever the mean of the perturbed population is negative, then the sub-circuit without them will be unstable.

### 5.10 Response distribution to partial and non-homogeneous perturbations

We now consider the case in which each population receives a partial, but the intensity of the perturbation is different for each perturbed neuron, mimicking disorder in the ChR2 expression. More specifically we need to recompute the expressions in equations (S68, S69) for the case in which we have a perturbation vector *δh* = {*δh*_1_, *δh*_2_, · · ·, *δh_n_*} instead of having entries like 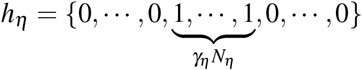, has entries given by 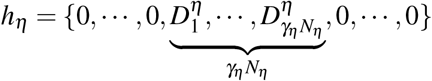, where 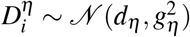.

The optogenetic targeting matrix Σ, instead of being given by Eq. (S99), will be in this case:

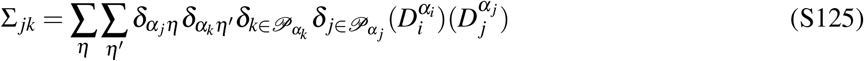

The expression for the perturbation to cell *i* will then be written as a mean given by the response that the network would have in the absence of disorder in the connectivity and a variance computed via Eqs (S68, S69). Specifically:

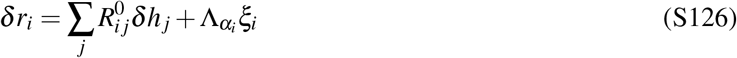

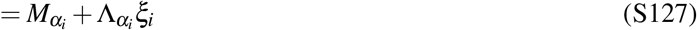

with

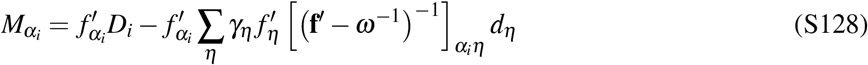

where 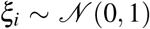 and 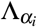 is the generalization of Eq. (S108) to disordered perturbations, obtained by replacing Eq. (S125) into (S98) (we note that the only term that needs to be re-computed is the term *M*).

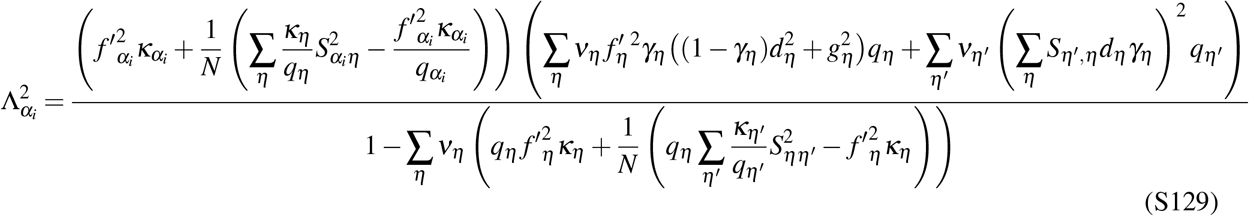

In the large N limit, this equation reduces to

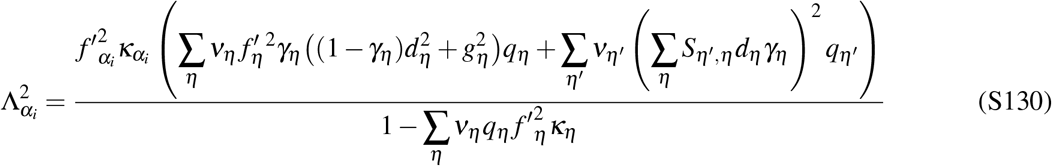

Which in the end means that the response of a neuron that belongs to the population *α_i_* will respond to the optogenetic perturbation with a mean and a variance given by

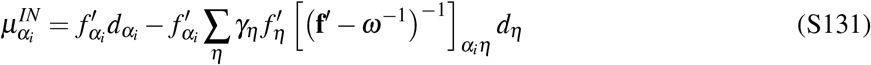

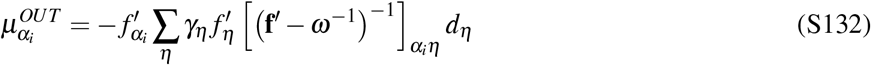

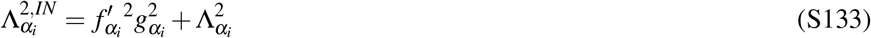

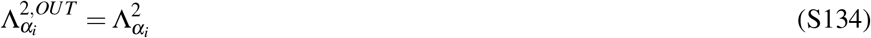

Analogously as before, we obtain a distribution of responses for the perturbed cells given by

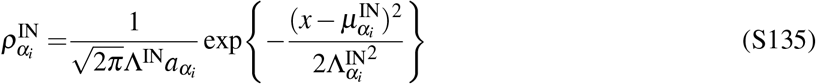

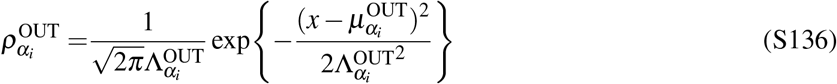

So the total distribution of responses is

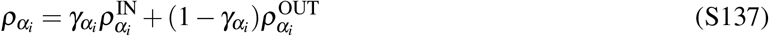

### 5.11 Link to the low-dimensional system linear response

The activity of the LD system is equivalent to the mean of the non-disordered high-dimensional system. Perturbing all the neurons in a population *α_j_* and then measuring the mean activity in the population *α_i_* should be equivalent to computing the linear response in the LD system. If the measuring vector 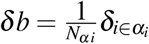 and *δh* is the optogenetic perturbation to all neurons in a given population, then

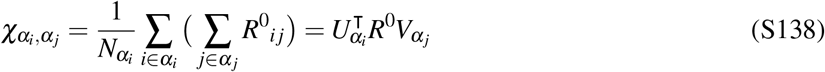

Inserting (S75) into the above expression we obtain:

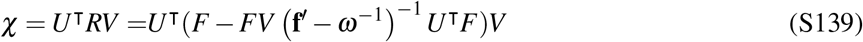

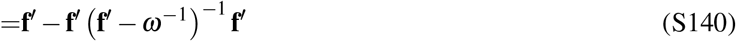

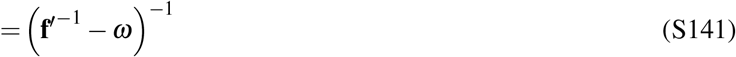

Which is the definition of *χ* as in Eq. (S9). We also need to show that the variance vanishes for large N. Writing 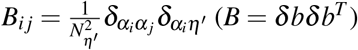, and inserting it and Eq. (S92) into Eq. (S69) we find that :

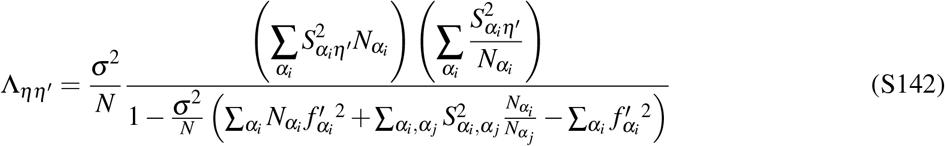

This variance vanishes for large N, making the usage of the small circuit as a limit of the average behavior of the large one rigorous for linear networks.

## 6 Eigenvalue Spectrum of the Jacobian in the HFA

In (Tao, 2013), it is shown that given a matrix with i.i.d. entries to which is added a deterministic matrix with rank 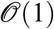 and operator norm 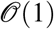, the distribution of eigenvalues of the Jacobian in the HFA is going to follow the circular law, except for a set of outlier eigenvalues. These outlier eigenvalues, are placed asymptotically in the same location as the eigenvalues of the low rank added matrix.

We now show that the Jacobian of our system under the HFA satisfies that condition. The scaling chosen for the low rank part of the connectivity matrix in the HFA guarantees that its operator norm is 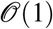. Using the definition of F and of **f**′ from Eq. (S61), the Jacobian under the HFA is written as

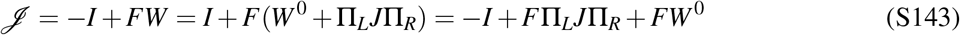

Let us for a moment write the non-disordered component of the connectivity as

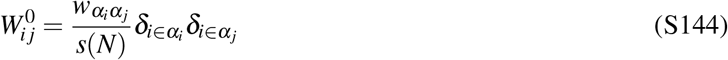

where *w_αβ_* is as before and s(N) is the scaling, which up to now we took to be *s*(*N*) = *N*. If we take U and V as defined in Eq.S72 and define the *n*-dimensional diagonal matrix *L* with elements *L_αα_* = *q^α^* = *N^α^ / N* and *I_n_* as the *n*-dimensional identity, then we have the following relations :

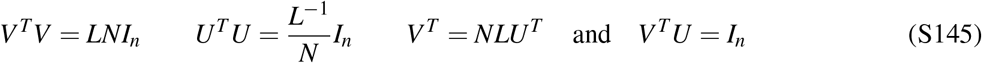

We want to compute the operator norm of the low-rank part 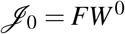, given by 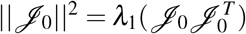 where *λ*_1_ is the max eigenvalue. We notice that 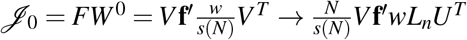. We notice that *ω* = *wL_n_* is the connectivity of the non-disordered LD system, and that therefore if *s*(*N*) = *N* then the nonzero eigenvalues of *J*_0_ are those of the LD system. In order to compute the eigenvalues of 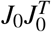 we notice:

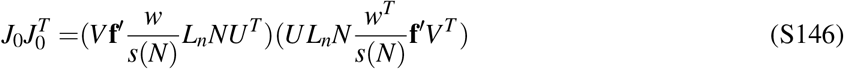

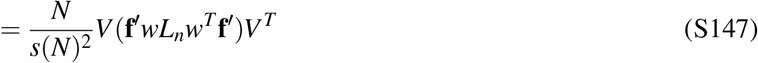

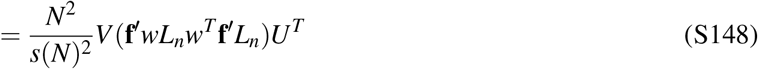

Where we used Eq S145 twice. This matrix then, has (N-n) zero eigenvalues and *n* eigenvalues that are those of 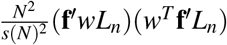. We notice that only if *s*(*N*) = *N* these eigenvalues will be 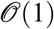. In the simplest case in which *F*Π_*L*_*J*Π_*R*_ is i.i.d., taking *s*(*N*) = *N* would be the only requirement to prove that the HFA has outlier eigenvalues if the LD system. For the case studied here, in which the disordered part of the connectivity is not *i.i.d*. but has a block-structured variance, there is not current proof that the deterministic outliers are those of the low-rank perturbation, but we nevertheless observe good agreement numerically.

## 7 Numerics

All numerical simulations and analyses were done with custom code in Python.

### 7.1 Network simulations

All network simulations were done with an Euler scheme with d=0.05 ms. The networks had N=3000 neurons with the number of E, PV, SOM and VIP cells being the 0.8 %,0.25 %,0.125 %, and 0.125 % of the total, respectively. We observe that due to the scaling in the model, none of these simulations depend on network size. Simulations usually ran for half a second (500 ms) each. All the simulations were done on models obtained as described in Sec. 4.4, with the additional requirement that they are stable, i.e. that the Jacobian of the network had all negative eigenvalues.

### 7.2 Linear response

The numerical calculation of the linear response was done by simulating the network, defining the matrix F as in Eq. S61 and numerically evaluating Eq. S62. Usually, 25 network instantiations were used to compute the distributions shown in the main text, given the relative small number of cells in the interneuronal populations.

### 7.3 Inference of locomotion inputs

In order to infer the locomotion inputs, fitted the distribution of changes in activity induced by locomotion per cell type and stimulus condition with a Gaussian distribution. This gave us 4 mean values and 4 variance values per contrast condition. This would correspond to the left hand side of Eq. S128, and S130 respectively. Now the only unknowns in those equations are *d_η_* (*c*) and *g_η_* (*c*) the (contrast dependent) locomotory inputs per cell-type *η*. These functions were chosen to be *d_η_* (*c*) = *aη* + *bη C*(*c*) where 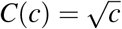 in the main text (Fig. 5) and *C*(*c*) = *c* or *C*(*c*) = 1 (linear with, or independent of, contrast) in the supplement (Fig. S5). By numerically solving for all the 8 constraints to occur for each contrast, we find the locomotion dependent inputs.

## Supplementary Figures

**Figure S1.**
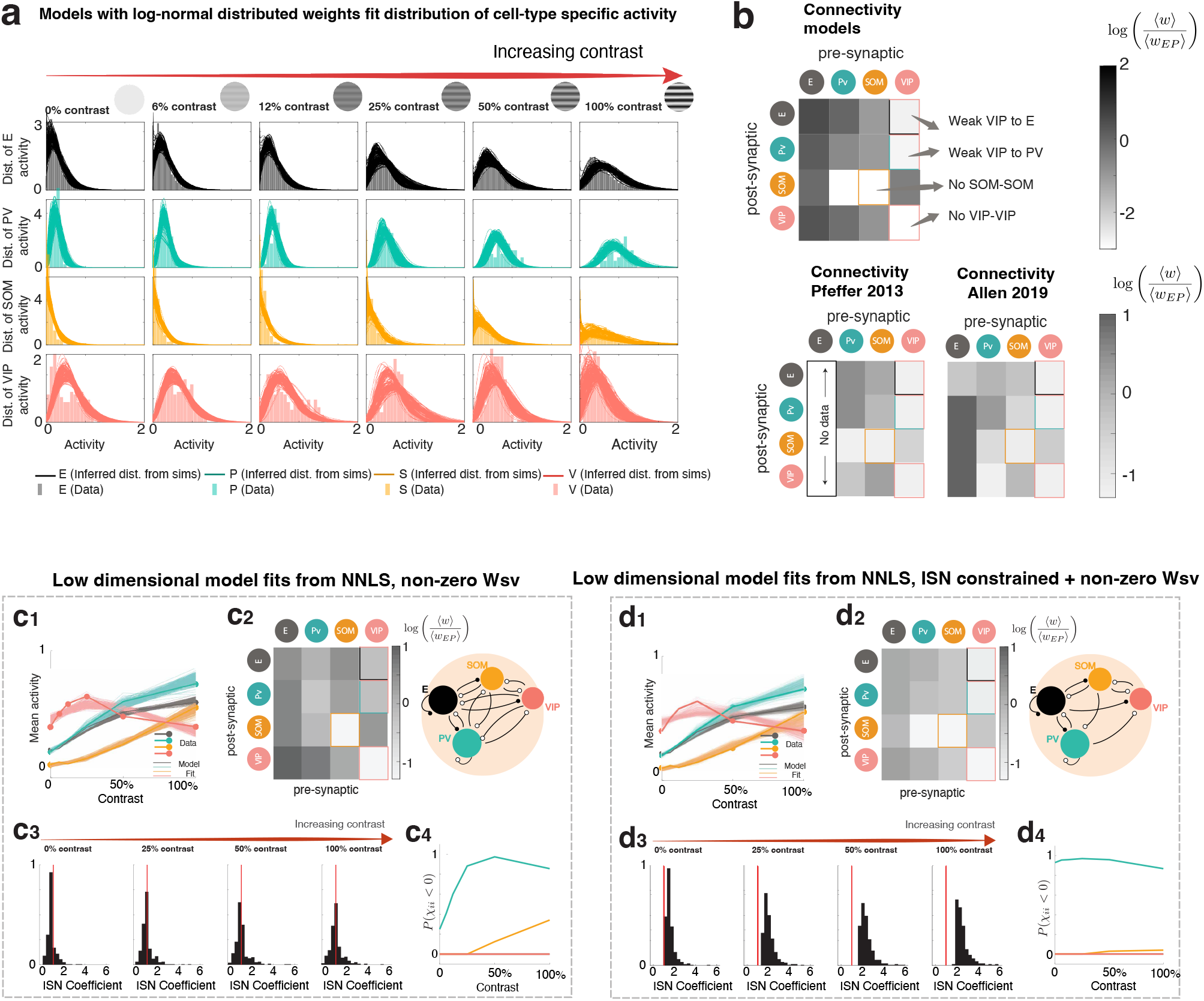
a) Large-scale models with log-normally distributed weights. The mean field theory used here is valid for any weight distribution whose tails decay as fast or faster than a Gaussian (Jonathan Touboul, personal communication). In particular any truncated distribution will obey this constraint. Here, we instead show that if we take the parameters found in the family of models and use non-constrained log-normal weight distributions, the models are still in good agreement with the data. Shown here are the extracted envelope (i.e. fitting the simulation like the data to extract the envelope given by Eq. S47) of simulations with log-normal weights for all models whose mean field fit has a total error smaller than 0.5. **b) Direct comparison with experimentally reported synaptic connectivity**. Top: Mean of the distributions of weights shown in 1, normalized by the synaptic weight from PV to E, for comparison with the available experimental data (background grayscale in panel 1**e**). Bottom Left: Synaptic weight connectivity as obtained in (Pfeffer et al., 2013). Bottom Right: Publicly available connectivity data from the Allen institute (https://portal.brain-map.org). The shown matrix is mean synaptic weight of the distribution of connections times the connection probability times the fraction of neurons belonging to the pre-synaptic cell type, normalized by the synaptic weight from PV to E, as done originally in (Pfeffer et al., 2013). These two matrices are shown here for comparison. There is currently no agreement on the strength of the connection from PV to VIP.

**Figure S2.**
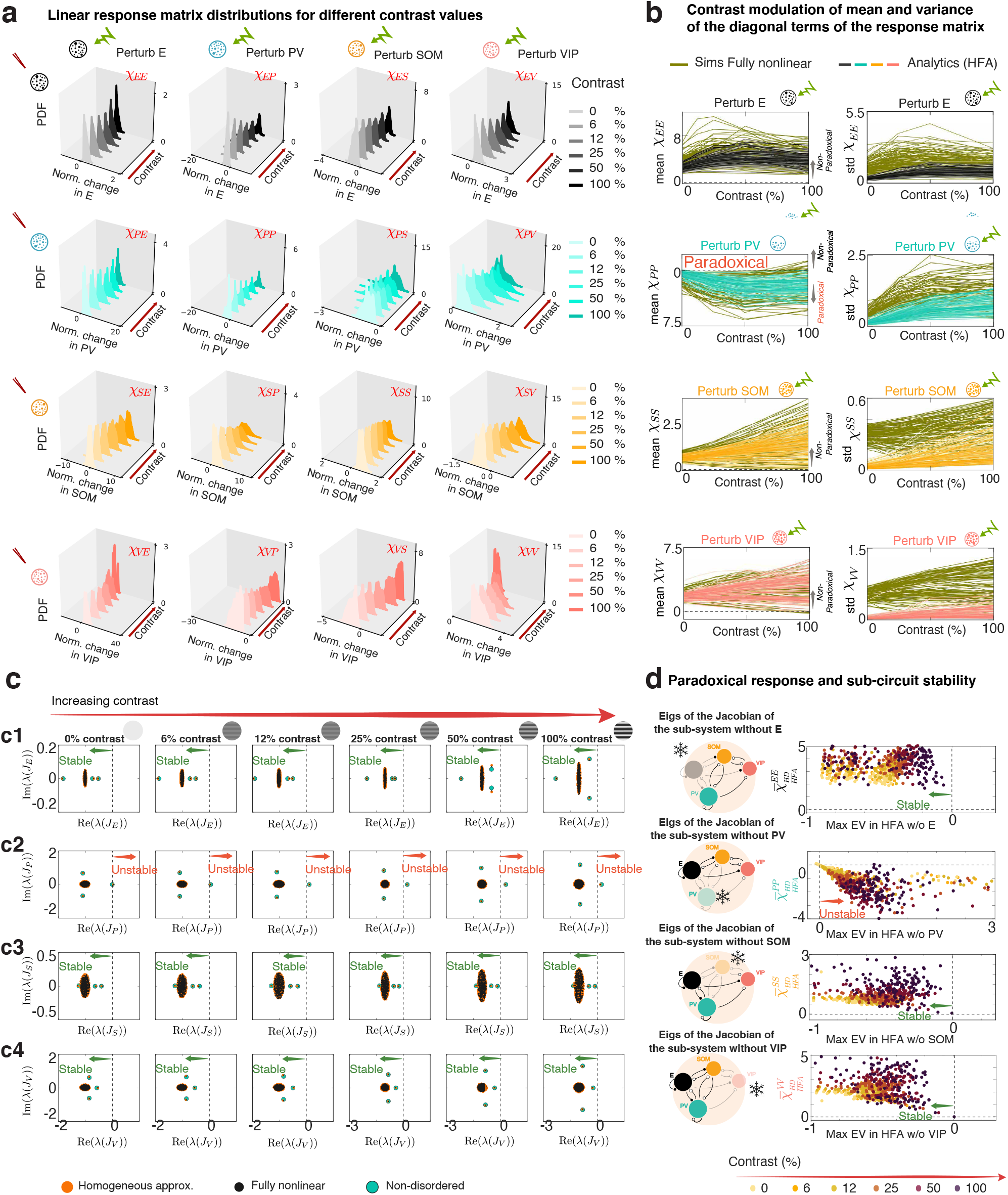
a) Response distributions to full population perturbations for different values of contrast in large-scale models. Distribution of linear responses to homogeneous full-population perturbations for different values of the contrast for all those models chosen in Fig. 1. **b)** Left panels: Mean response of each cell type to a perturbation to that same cell type as a function of contrast, for models in the HFA (E in black, PV in turquoise, SOM in orange, VIP in pink) and for the fully nonlinear network (green). Right panels: Standard deviation of the response. Note the paradoxical response of PV at all contrasts for almost all models and the non-paradoxical response of SOM in most cases. In the multiple-cell-type circuit and unlike in the EI system, excitatory activity can in principle also respond paradoxically (Miller and Palmigiano, 2020). Nevertheless, none of the data-compatible models obtained had an excitatory paradoxical mean response.Mean response of each cell type to a perturbation to that same cell type, vs the real part of the maximum eigenvalue of the sub-circuit without that cell type, for all values of the contrast. **c) Linking stability and response to full population perturbations in large-scale models. c**_1_)Eigenvalues of the Jacobian of the sub-circuit without the entire E population (i.e. *J_E_*) as a function of contrast. The response of pyramidal cells is never paradoxical and the sub-circuit without it is always stable, both in the system using the HFA approximation (orange), the non-disordered system (i.e. the LD system, turquoise), and the fully-non linear system without approximations (black). **c**_2_) Idem c_1_) for the sub-circuit without PV (*J_P_*). The sub-system without PV is always unstable, with only one eigenvalue positive, guaranteeing a paradoxical response (see also Miller and Palmigiano, 2020). **c**_3_) for the sub-circuit without SOM (*J_S_*), although this sub-circuit approaches instability for large contrast in this particular example it remains non-paradoxical. **c**_4_) for the sub-circuit without VIP (*J_V_*). **d)** Mean response of each cell type to a perturbation to that same cell type, vs the real part of the maximum eigenvalue of the sub-circuit without that cell type, for all values of the contrast.

**Figure S3.**
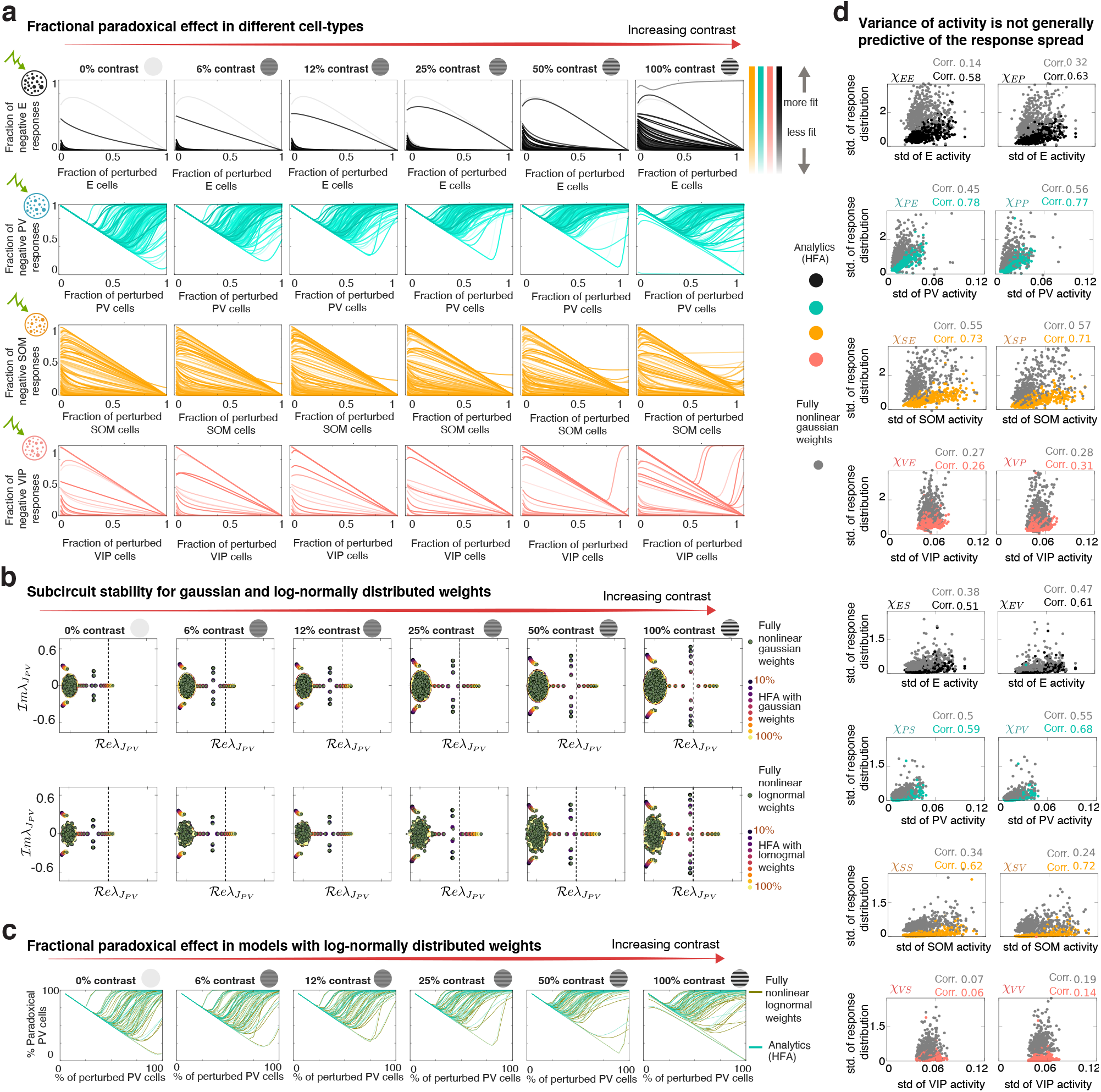
a) Fractional paradoxical effect in the family of large-scale models with the HFA. Dependence of the fraction of negative responses of the E (top), PV (middle top), SOM (middle bottom) and VIP (bottom) cells as a function of the fraction of stimulated cells in the same population. Shaded color indicates fitness. Notice how few models have a VIP fractional paradoxical effect in VIP at high contrast. **b) Linking stability and response to partial perturbations in large-scale models**. (Top) Eigenvalue distribution of the sub-circuit obtained by ‘freezing’ or removing different percentages of PV neurons from it, for an example model with Gaussian distributed weights. Different colors indicate different fractions of removed PV neurons in the HFA network (dark blue without 10% of PV neurons, yellow without the entire PV population). Dots in green are the corresponding eigenvalues of the sub-circuit in the fully nonlinear case. The eigenvalue distribution in the HFA and the fully nonlinear case match perfectly. (Bottom) for the same example network but with log-normally distributed weights. **c) Fractional paradoxical response for networks with log-normally distributed weights**. The models show a fractional paradoxical effect which is in good agreement to the effect described in networks with HFA with Gaussian weights, for which the theory of response to perturbations is exact. **d) Spread of the activity distribution is a poor predictor of the spread of optogenetic responses**. Standard deviation of the responses of each cell type to perturbations to each cell type as a function of the standard deviation of the activity, for all values of contrast. We see that the correlation between the spread of these distributions is only mild, both under the analytical approximation (black E, turquoise PV, orange SOM and pink VIP), and for the fully nonlinear system in grey. The spread of the distribution of activity is therefore not a strong predictor of the distribution of responses

**Figure S4.**
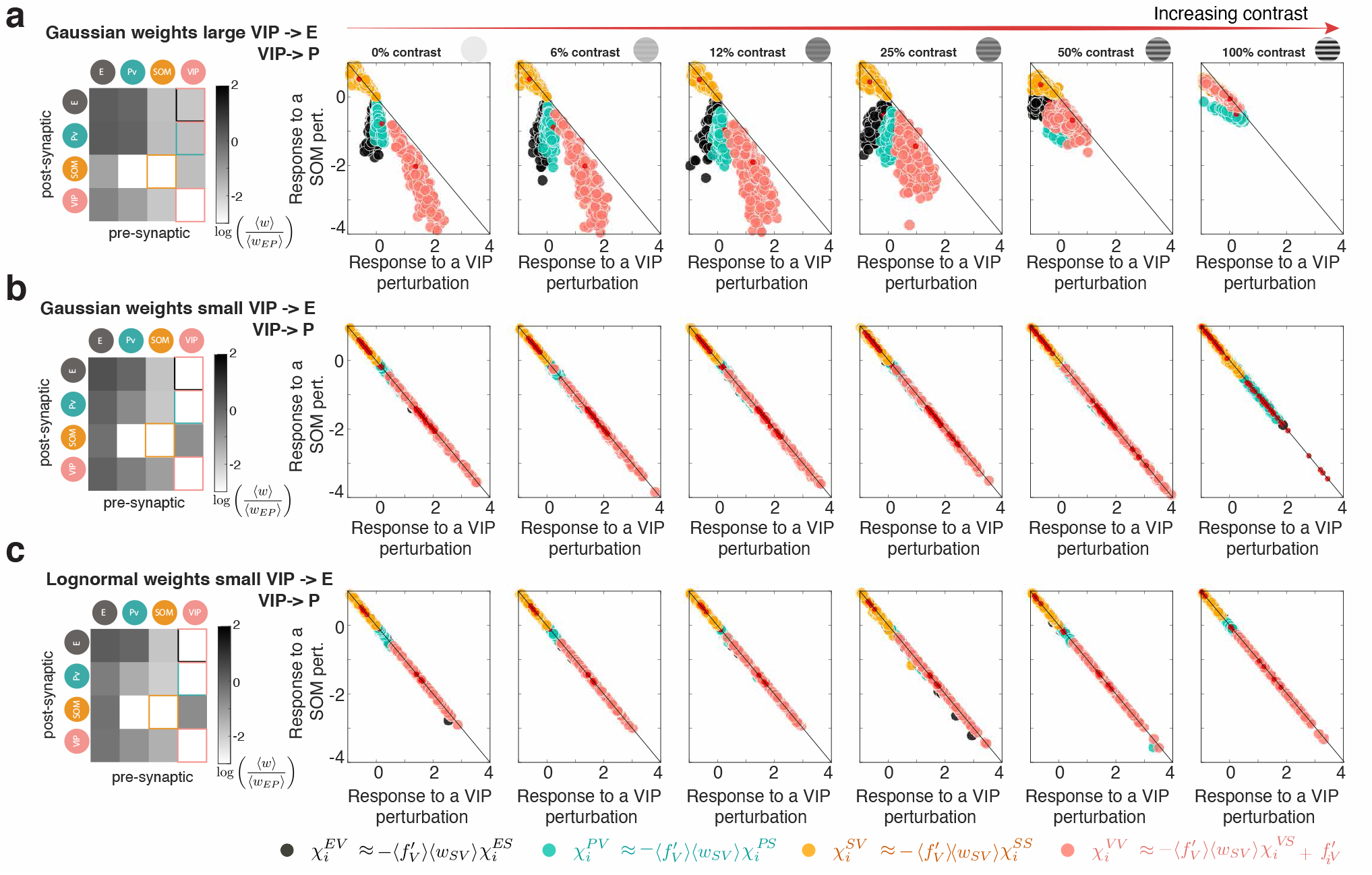
Disinhibition symmetries in the response only depends on the VIP projections and not the shape of the weight distribution. **a)** Left: Mean connectivity of a model in which the mean projections from VIP to E (*w_EV_*) and to PV (*w_PV_*) are large. Right: Response of each cell to a full SOM perturbation vs the response to a full VIP population, In this case, the LD hidden symmetries are not respected (red points, as in Fig. 3, are outside the negative diagonal). Furthermore, the entire distribution of responses to a VIP and a SOM perturbation does not obey a contrast-independent relation. Still, there is a clear correlation between the response to a SOM and a VIP perturbation for each cell. **b)** Model with KL error smaller than 0.5 and *w_EV_* > −0.1. Left: mean connectivity across models in this configuration. Right: All these models show clear HD symmetries. **c)** Top 10% of models with KL error smaller than 0.5 and *w_EV_* > −0.1, simulations done with a log-normal distribution of weights. Left: mean connectivity across models in this configuration. Right: All these models, even with heavy tailed weight distributions, show clear HD symmetries.

**Figure S5.**
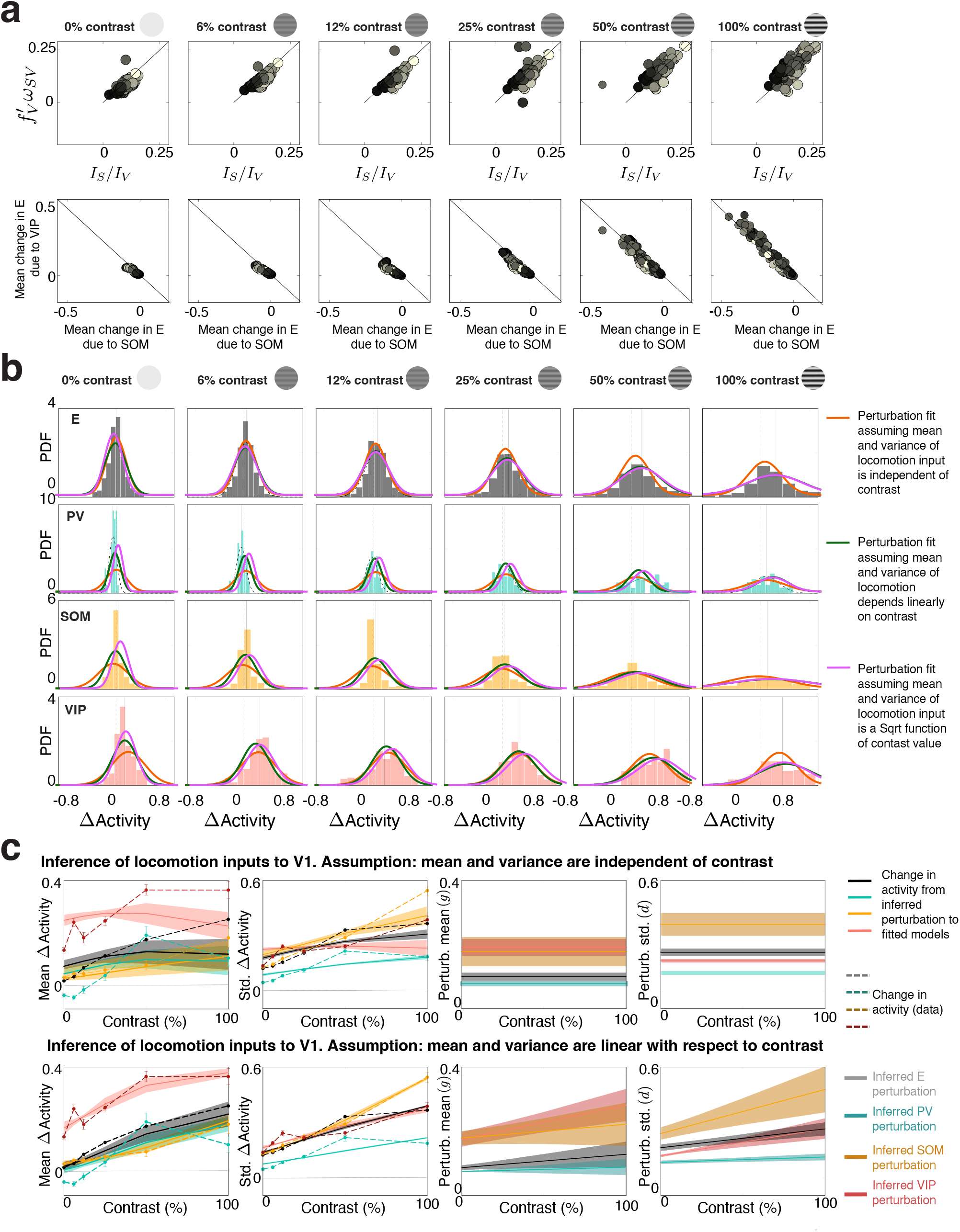
Strong and wide inputs to VIP and SOM lead to network responses that mimic the effect of locomotion. **a)** Top: Scatter plot showing that the disinhibition symmetries underlie behavioral modulations of activity. The theory predicts that, if the ratio of the inputs to SOM and VIP are proportional to the mean VIP gain (under the HFA), then the net effect of those perturbations in E will indeed cancel (see Eq. S30). Here we show that when we infer the locomotory inputs to SOM and VIP, they are proportional to the mean VIP gain times the mean strength of the connection from VIP to SOM (different for each model). This is the case even when the stimulus contrast is large and the mean change in E due to locomotion no longer vanishes (see below). Bottom: Change in E activity due locomotory inputs to SOM Vs due to locomotory inputs to VIP. At the maximum contrast, models largely lie on the diagonal and only few models show disinhibition (i.e inputs lead to an increase in E activity). **b)** Distribution of Δ activity (difference between each cell’s activity in the locomotion and the stationary condition) for each cell type and different values of the stimulus contrast. Dashed lines indicate best Gaussian fits. Solid lines are fits from the explicit expressions (see Eqs. S126 and S129) under three different assumptions i) mean and variance of the inputs to each population are independent on contrast (orange, corresponding to panel b) ii) mean and variance of the inputs to each population depend linearly on contrast (green, corresponding to panel c) and ii) mean and variance of the inputs depend as the square root of the contrast (magenta, same curves as in main text. **c** Top: Mean of Δ activity and standard deviation of Δ activity as a function of contrast. Dashed lines are the data (E in black, PV in blue, SOM in orange and VIP in dark red) and full lines are the mean of the distribution of fits as described in **a**, for the family of models that fit the stationary data, under the assumption that locomotory inputs do not depend on contrast. The mean (g) and the standard deviation (d) of the inferred perturbation are shown in the right panels, and independent of stimulus properties. Bottom: Same as top but requiring the locomotory inputs to depend linearly on the contrast.

